# Novel antiproliferative tripeptides block AP-1 transcriptional complex by in silico approach

**DOI:** 10.1101/2020.05.08.083972

**Authors:** Ajay Kumar Raj, Jainish Kothari, Sethamma TN Sinchana, Kiran Lokhande, K. V. Swamy, Nilesh Kumar Sharma

**Affiliations:** Cancer and Translational Research Lab, Dr. D.Y. Patil Biotechnology & Bioinformatics Institute, Dr. D.Y. Patil Vidyapeeth, Pune, Maharashtra, India, 411033

**Keywords:** Microbiomes, Metabolites, Tri-peptides Cancer Therapy, Mimic, Transcription factor, down regulation of gene expression

## Abstract

**BACKGROUND:** The complexity and heterogeneity at genetic, epigenetic and microenvironment levels are key attributes of tumors. Genetic heterogeneity encompasses one of key factors at transcriptional gene regulation that promote abnormal proliferation, invasiveness and metastasis. Among various key pro-tumor transcriptional complexes, activating protein-1 (AP-1) transcriptional complex controls the transcriptional expression of key oncogenes in cancer cells. Therefore, an avenue to search for a chemical inhibition approach of the AP-1 transcriptional complex is warranted in cancer therapeutics.

**METHODS:** To achieve chemical inhibition of AP-1 transcriptional complex, we report novel tripeptides identified from the goat urine DMSO fraction as potential agents that bind to AP-1 responsive TPA element and heterodimer c-Jun:c-Fos. Novel tripeptides enriched GUDF were tested against DNA substrates to assess DNA metabolizing activity. Further, Novel tripeptides enriched GUDF were treated upon HCT-116 cells to estimate the nature of tripeptides entered into the intracellular compartment of HCT-116 cells. Here, we report on a novel methodology that employ VTGE assisted intracellular metabolite purification and is analyzed with the help of LC-HRMS technique. Post purification of intracellular metabolites that included tripeptides of GUDF, these tripeptides from DMSO and GUDF treated HCT-116 cells were subjected to molecular docking and ligand-DNA:AP-1 (PDB ID: 1FOS) interaction study by using bioinformatics tools AutoDock Vina and PyMol.

**RESULTS:** GUDF enriched with tripeptides and other metabolites show appreciable instability of DNA substrates plasmid and genomic DNA to an extent of 90%. Interestingly, LC-HRMS analysis of intracellular metabolite profiling of GUDF treated HCT-116 cells reveal the appreciable abundance of tripeptides Glu-Glu-Arg, Gly-Arg-Pro, Gln-Lys-Arg, Glu-Glu-Lys, Trp-Trp-Val. On the other hand, DMSO treated HCT-116 cells show the presence of Ser-Trp-Lys, Glu-Glu-Gln, Glu-Glu-Lys, Ser-Leu-Ser. Interestingly, GUDF treated HCT-116 cells show inhibition of proliferation by more than 70%. Among the identified intracellular tripeptides, Glu-Glu-Arg (9.1 Kcal/Mol), Gly-Arg-Pro (8.8 Kcal/Mol), and Gln-Lys-Arg (6.8) show a precise and strong binding to heptameric TPA response element 5’ TGAGTCA 3’ and key amino acid residue within the AP-1 transcriptional complex.

**CONCLUSION:** In summary, this study suggests the potential of novel tripeptides, those are reported from GUDF intracellularly in HCT-116 cells to destabilize the AP-1 transcriptional complex. Data indicate that cellular arrest in HCT-116 cells treated by GUDF is well supported by the molecular docking observations that destabilization of AP-1 complex is linked to reduced growth and proliferation.

## INTRODUCTION

Cancer is one of the most prevalent pandemics all over the world causing millions of deaths every year (1-2). At global level, lung cancer, breast cancer and colon cancer are leading cancer types in terms of incidences and mortality rate. Besides conventional anticancer therapy, there are therapeutic avenues to target molecular distinctiveness within the tumors including genetic, transcriptional and microenvironmental heterogeneity (3-9).

In essence, genetic and transcriptional heterogeneity are one of key factors that maintain the pro-tumor microenvironment (10-13). Among various driving factors as oncoproteins, transcription factors are large proportions of proteins that bind DNA helix at specific regulatory response elements in order to modulate the expression of a set of genes that contribute towards distinctive features of cancer cells including growth and proliferation (14-18).

Transcription factors are known to achieve gain of function that allow the cancer cells to allow the abnormal expression of a set of genes dedicated to hallmarks of tumor including uncontrolled proliferation (3-6, 19-25). In a pool of oncogenic transcription factors, AP-1 transcription factor complex consists of c-Fos and c-Jun heterodimers having similar sequence and structure with a bZip protein which helps the AP-1 complex to bind upon specific heptamer consensus nucleotide sequence 5’TGAGTCA3’ (AP-1 site) (3, 10-13). This heptamer is also referred to as the 12-O-Tetradecanoylphorbol-13-acetate (TPA) response element.

In recent, blockade of AP-1 transcriptional complex is perceived as a promising option to reduce the uncontrollable growth and proliferation of cancer cells by using small molecular inhibitors and peptide mimetics (14-25). These small molecule inhibitors are suggested to bind precisely with the TPA response element of target genes by non-covalent forces such as the coulombic force, vander-waals interactions, and hydrogen bonding and that may be responsible for their anti-cancer properties. (19-25). Evaluation of these molecular inhibitors are evaluated by various approaches including in vitro transcriptional assay and molecular docking computational approach (15-26).

There are limited findings that support the use of small molecular inhibitor such as nor-dihydroguaiaretic acid (NDGA), curcumin and tripeptides against the key oncogenic transcriptional complex such as AP-1, FOXO1, MYC and hypoxia inducing factor-3 (14-28). In fact, chemical inhibition of AP-1 transcriptional complex has shown encouraging evidence on retardation of proliferative potential of cancer cells during in vitro and in vivo experiments (29-37).

In search of potential small molecular inhibitors and tripeptides as anticancer drugs, limited data indicate that biological fluids such as plasma, urine, milk from human and ruminants contains tripeptides and other metabolites that are shown to display biological properties such as DNA destabilizing and chelation activity, anti-inflammatory and modulation of cellular growth and proliferation (38-46). Among various small molecules, tripeptides are suggested to be transported across the cell membrane by the help of peptide transporters, endocytosis and membrane permeation (47-54). We have also shown that goat urine derived metabolites and tripeptides enriched fraction displayed DNA metabolizing and anti-proliferative activity (45-46).

Based on the existing thrust to explore AP-1 transcriptional complex inhibitors, we propose to test the ability of novel tripeptides displaying anti-proliferative effects as a potential inhibitor of AP-1 transcriptional complex.

## MATERIALS AND METHODS

### Materials

Cell culture reagents were procured from Himedia India Pvt. Ltd. and Invitrogen India Pvt. Ltd. The HCT-116 cells were obtained from the National Centre For Cell Science (NCCS), Pune, India. Plasmid DNA pBR322, DMSO, acrylamide and other chemicals were of molecular biology grade and these chemicals were purchased from Himedia India Pvt. Ltd and Merck India Pvt. Ltd.

### Cell Line Maintenance

The HCT-116 cells were cultured and maintained in DMEM (Dulbecco’s Modified Eagles Medium) (Himedia) with high glucose supplemented with 10% heat-inactivated FBS/penicillin (100 units/ml)/streptomycin (100 μg/ml) at 37°C in a humidified 5% CO_2_ incubator. The passaging number of HCT-116 cells was maintained below twenty.

### *In Vitro* DNA metabolizing assay

Goat urine DMSO fraction (GUDF) enriched with tripeptides was used for the evaluation of DNA metabolizing effects by in vitro reaction. In brief, GUDF enriched with tripeptides is prepared in sterile DMSO and filtered by using 0.45 μm syringe driven filter and details are adopted from previously published methodology (45-47). An in vitro reaction constituted components including pBR322 (plasmid DNA) and genomic DNA (HCT-116 cells) of (100 ng/μl) that were added along with 2.5 μl TAE buffer (Tri-acetate/EDTA 10 mM, pH 7.4). This reaction mixture was treated by different concentration 0.1 μl, 0.5 μl and 1 μl of GUDF (Stock concentration 10 mg/ml). Finally, total volume of reaction was brought to 25 μl by the addition of nuclease free water. Next, incubation of DNA metabolizing reaction was permitted at 37°C for 1 hr. Finally, reaction was terminated and a standard protocol for DNA electrophoresis, visualization and densitometry was adopted and performed (45-47).

### Trypan blue dye exclusion assay

HCT-116 cells were plated onto six well plate (15×10^4^ cells per well) and incubated in the presence of complete fresh DMEM high glucose medium supplemented with 10% FBS. After 16-18 h, cells were treated with 2 ml of complete fresh DMEM medium containing 5 μl and 10 μl GUDF (stock concentration 10 mg per ml) with 25 and 50 μg/ml final concentration. DMSO solvent with the same volume was treated upon HCT-116 cells to serve as a negative. At the end of 72 hr of incubation, a standard protocol was adopted to assess the presence of a total and viable HCT-116 cell treated by GUDF and DMSO (46).

### Propidium iodide/Annexin V staining assay

HCT-116 cells were seeded in duplicates into six well plate at the plating density of 15×10^4^ cell per well. After 16-18 hours of plating, HCT-116 cells were treated with10 μl GUDF (stock concentration 10 mg per ml) achieving 50 μg/ml final concentration and DMSO for 72 hr. At the end of treatment, HCT-116 cells were harvested by standard protocol. In brief, Annexin V/FITC apoptosis detection kit (ThermoFisher, USA) was used to stain HCT-116 cells for the assessment of viability and apoptosis. Here, flow cytometry analysis was performed on BD FACSJazz Cytometer. The detailed methodology during flow cytometry was adopted from previously reported methods (47, 53).

### Purification of intracellular metabolites by VTGE

To study the intracellular metabolites in HCT-116 cells treated with GUDF and DMSO, we prepared hypotonically lysed whole cell lysate of HCT-116 cells as adopted from previously reported protocol (53). In brief, for 10 million harvested HCT-116 cells, we used 500 μl of hypotonic lysis buffer (10 mM KCl, 10 mM NaCl, 20 mM Tris, pH 7.4). Further, sterile membrane filtered clear hypotonically lysed whole cell lysates of HCT-116 (250 μl) cells were diluted to 750 μl by the addition of μl 500 hypotonic buffer. In brief, preparation of hypotonically lysed whole cell lysate of HCT-116 cells was adopted from previously reported protocol (53). To achieve the purification of intracellular metabolites, above prepared HCT-116 cell lysate (750 μl) was added with 250 μl of 4X loading buffer (0.5 M Tris, pH 6.8 and Glycerol). Next, these sample mixtures were loaded and allowed to get separated on a specifically designed VTGE system (Figure 1). Precisely, we used 15% acrylamide gel (acrylamide:bisacrylamide, 30:1) as a gel matrix that helped to trap high M.W proteins in the gel and low M.W. metabolites including tripeptides were eluted in the elution buffer. The detailed specifications and working conditions of VTGE were adopted from previously reported methodologies (46, 53).

**Figure 1.**
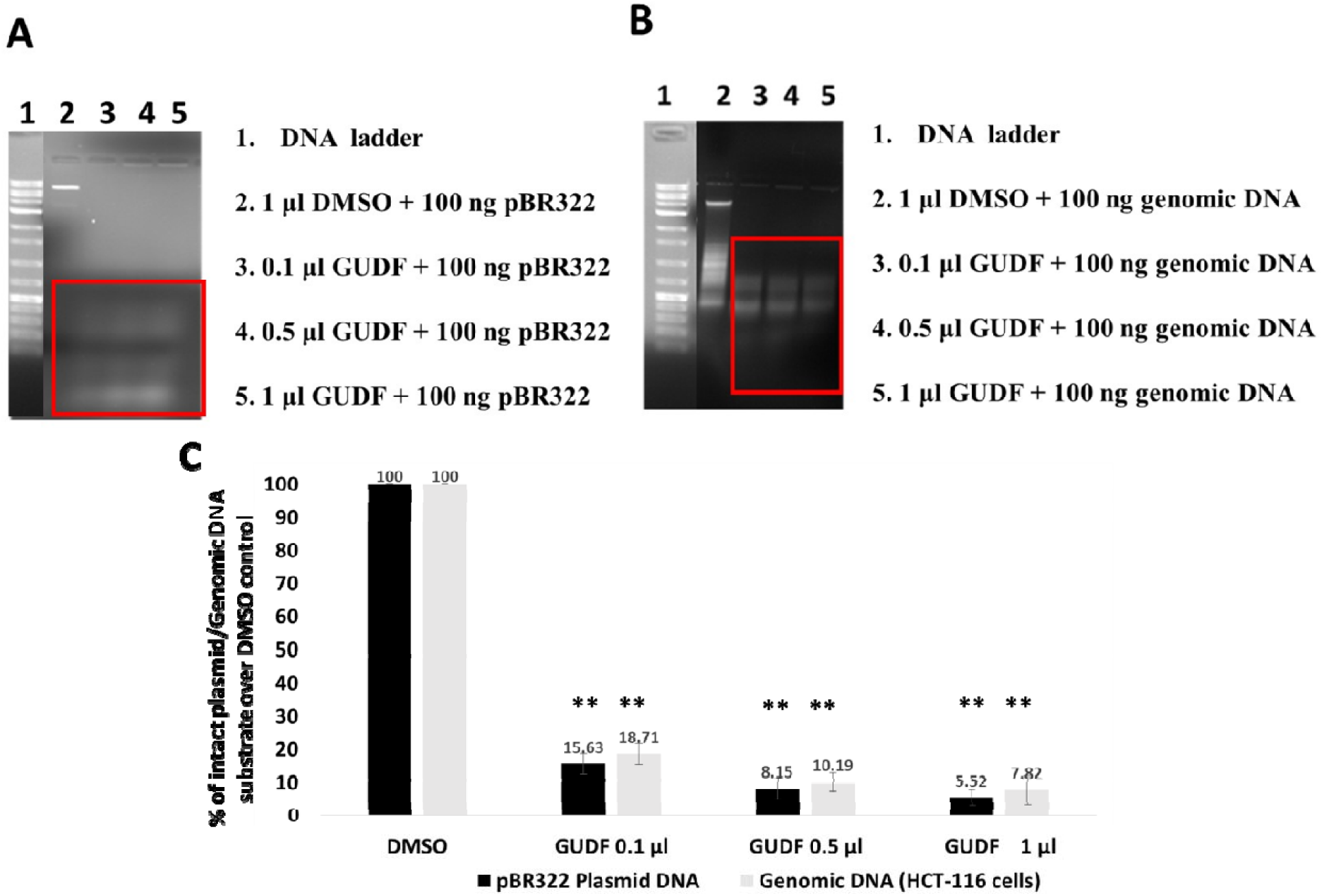
GUDF enriched with tripeptides show DNA metabolizing activity. (A). This photograph shows an ethidium bromide stained agarose gel electrophoresis-based separation of in vitro plasmid DNA pBR322 treated by varied concentration of GUDF. (B) This photograph represents an ethidium bromide stained agarose gel electrophoresis-based separation of genomic DNA treated by varied concentration of GUDF. The image was visualized and captured using BIO-RAD EZ imaging system. (C) This bar graph show the reduction in th intact plasmid DNA and genomic DNA treated with GUDF over DMSO control. DMSO control was used as solvent control. The image was visualized and captured using BIO-RAD EZ imaging system. Data are represented as mean ± SD. Each experiment was conducted independently three times. The bar graph without an asterisk denotes that there is no any significant difference compared to DMSO control. * Significantly different from DMSO control at the P-value < 0.05. ** Significantly different from DMSO control at P-value < 0.01.

### Intracellular tripeptide metabolite profiling by LC-HRMS

Further, the eluted intracellular metabolite of HCT-116 cells was taken for LC-HRMS to know about the metabolites including tripeptides accumulated inside the cells. In brief, a platform of Agilent TOF/Q-TOF Mass Spectrometer station equipped with an ion source Dual AJS ESI was employed for MS/MS analysis. For liquid chromatography, separation of purified intracellular metabolites was achieved by using RPC18 Hypersil GOLD C18 100 x 2.1 mm-3 μm column. Further, MS/MS acquisition of data were performed in positive and negative ionization mode. During acquisition and analysis of sample, methods and processes were adopted from previously reported methodologies (46,53).

### Molecular Docking Study on tripeptides and AP-1 transcriptional complex

The structures of novel tripeptides identified in the intracellular compartment of HCT-116 cells are detailed as Glu-Glu-Arg (CHEBI ID:144557), Gly-Arg-Pro (CHEBI ID:144473), Gln-Lys-Arg (CHEBI ID:144723), Glu-Glu-Lys (CHEBI ID:144461), Trp-Trp-Val (CHEBI ID:144555), Ser-Trp-Lys (CHEBI ID:144474), Glu-Glu-Gln (CHEBI ID:144559) and Ser-Leu-Ser (CHEBI ID: 144475). Besides these tested tripeptides, this study included earlier reported compounds NDGA (CHEBI ID: 7625), ethidium bromide (CHEBI ID: 4883), curcumin (CHEBI ID: 3962) and Hoechst 33342 (CHEBI ID: 51232) were downloaded in SDF format from the ChEBI database (https://www.ebi.ac.uk/chebi/). Then these tripeptides and compounds were converted using OpenBable into PDB format assigned with 3D coordinates. The target of Fos-Jun-DNA complex was downloaded directly from Protein Data Bank (PDB ID-1FOS) (https://www.rcsb.org/) and no further modifications were done to the crystal structure of Fos-Jun-DNA complex (10). The chains of ProA-ProB-DNAA-DNAB were taken for the *in-silico* study on AutoDock Tool 4.2.1 (ADT). We removed all the water molecules and added Kollman charges, assigned AD4 charges and added polar interactions using H-bonds. In this paper, to achieve molecular docking between tripeptides and other compounds as an inhibitor of AP-1 transcriptional complex, AutoDock Vina, an improvement with speed and accuracy is used (26). AutoDock Vina is known to calculate automatically grid maps. Furthermore, AutoDock Vina provides better clustering of results in the perspectives of unbiased data to the user. All tripeptides were then docked into the binding site of the receptor to predict binding conformations for the tripeptides. Here, blind docking was applied for all docking experiments that employed a grid box that was sufficient enough to encompass a potential ligand receptor complex (c-Jun:c-Fos:DNA). Here, grid box configurations including center_x = 54.694, center_y = 2.833, center_z = −19.441, size_x = 62, size_y = 76, size_z = 64 and energy_range = 4 are applied for all ligand-1FOS molecular interaction studies. The best conformation was determined by the binding affinity of tripeptides with the c-Jun:c-Fos:DNA Complex. Furthermore, binding position of tripeptides and other known compounds with potential to bind to TPA response element, c-Jun, c-Fos within the 1FOS as a part of docking structure was rendered by PyMol (www.pymol.org) view interaction studies.

## STATISTICAL ANALYSIS

Data are presented as the mean ± SD of at least three independent experiments. Differences are considered statistically significant at P < 0.05, using a Student’s t-test.

## RESULTS AND DISCUSSION

### Biological activities of GUDF enriched with tripeptides

In view of enrichment of biological fluids such as urine and plasma, there are reports on abundance of biologically active compounds such as organic acids and tripeptides (38-46). However, limited attempts have shown the potential of these tripeptides with DNA metabolizing and modulatory effects on growth and proliferations of normal and cancer cells.

Therefore, by employing a sterile and standard approach to fractionate active components enriched with tripeptides and their biological effects are reported earlier (45-46). In this paper, we looked into the detailed pattern of DNA metabolizing activity of GUDF enriched with tripeptides upon pBR322 and genomic DNA as substrates. In this simple assay, DNA instability effects of GUDF enriched with tripeptides is very clear and more than 90% of plasmid DNA and genomic DNA are fragmented (Figure 1A, B and C). Here, authors would like to draw attention that the pattern of damage to DNA is not in the form of smear of DNA that is mostly detected with contaminated samples with heavy metals and external contaminations. Therefore, these data encouraged us to suggest that used GUDF enriched with tripeptides samples are free from interfering components. Additionally, we have earlier shown that by autoclaving of GUDF enriched with tripeptides, the DNA metabolizing effects are absent and this is possibly due to the instability of organic nature of contributing agents in the form of tripeptides. Here, our observations are not surprising, because there are limited reports that suggest the selected tripeptides show the DNA metabolizing and binding activities (27-28, 47-54).

Furthermore, we proposed to collect molecular and cellular evidence that may be contributed by GUDF enriched with tripeptides along with DNA metabolizing effects. In this direction, sterile GUDF enriched with tripeptides were tested for growth and proliferation modulatory effects on HCT-116 cells. The selection of HCT-116 cells was based on the previous screening of GUDF enriched with tripeptides against various cancer cell lines including breast, colorectal and cervical cancer. Here, the intent of using GUDF enriched with tripeptides is not only for the evaluation of anti-growth and proliferation potentials, but also to explore the evidence on the nature of tripeptides that may be able to enter into the treated cancer cells. Data indicate the significant reduction in the HCT-116 cells treated by GUDF enriched with tripeptides that was revealed by the simple Trypan blue dye exclusion assay. Here, data hinted that GUDF enriched with tripeptides showed the clear inhibition of proliferation up to 28.97% and simultaneously reduction of viable cells are observed up to 16.54% (Figure 2A, B) during 72 hr of treatment. Surprisingly, data do not support the ability of GUDF enriched with tripeptides to cause the loss of viability of HCT-116 cells. But GUDF is able to arrest the proliferation of HCT-116 cells. Furthermore, data analysis by PI/Annexin V staining assay support the observations of Trypan dye exclusion assay that GUDF enriched with tripeptides do not show the significance presence of clear apoptotic death of HCT-116 cells, instead of up to 58.95% of cells are in the early apoptotic stage during proliferation arrest for a long period of treatment (Figure 2 C, D). There are views that treatment by drugs for long period of time may force cancer cells to the early apoptotic stage in case of the ability of drugs to bring proliferation arrest. Therefore, data indicated us to believe that GUDF enriched with tripeptides show proliferation arrest of HCT-116 cells, but not clear apoptotic cell death. This observation led us to speculate that GUDF enriched with tripeptides do not have components that can cause death of HCT-116 cells, but possibilities of compounds in the form of tripeptides that may show the intracellular effects to achieve the proliferation arrest. Taken together, GUDF enriched with tripeptides show appreciable DNA metabolizing and arrest of cellular proliferation of HCT-116 cells. Our observations are in consonance with the previously reported natural and synthetic tripeptides that reported on DNA binding, DNA metabolizing and modulations of cellular growth and proliferations (47-54). Interestingly, some tripeptide seryl-histidine, Lys-Gly-His-derived metallopeptides, Lys-Trp-Lys, Gly-His-Lys, Phe-Phe-Phe, Arg-Gly-Gly, Gly-Ala-His, and Gly-His-Lys show DNA binding and cellular toxicity to cancer cells (27-36, 47-54). These reported tripeptides are basic in nature, therefore, an attempt to reveal the nature of GUDF tripeptides in HCT-116 cells may shed light on the mechanisms of cellular effects. Hence, observed proliferation arrest of HCT-116 cells by GUDF enriched with tripeptides is conceivable in line with the existing set of anticancer drugs. At the same time, the molecular basis of effects by GUDF enriched with tripeptides needs further investigations by in vitro and in-silico approaches.

**Figure 2.**
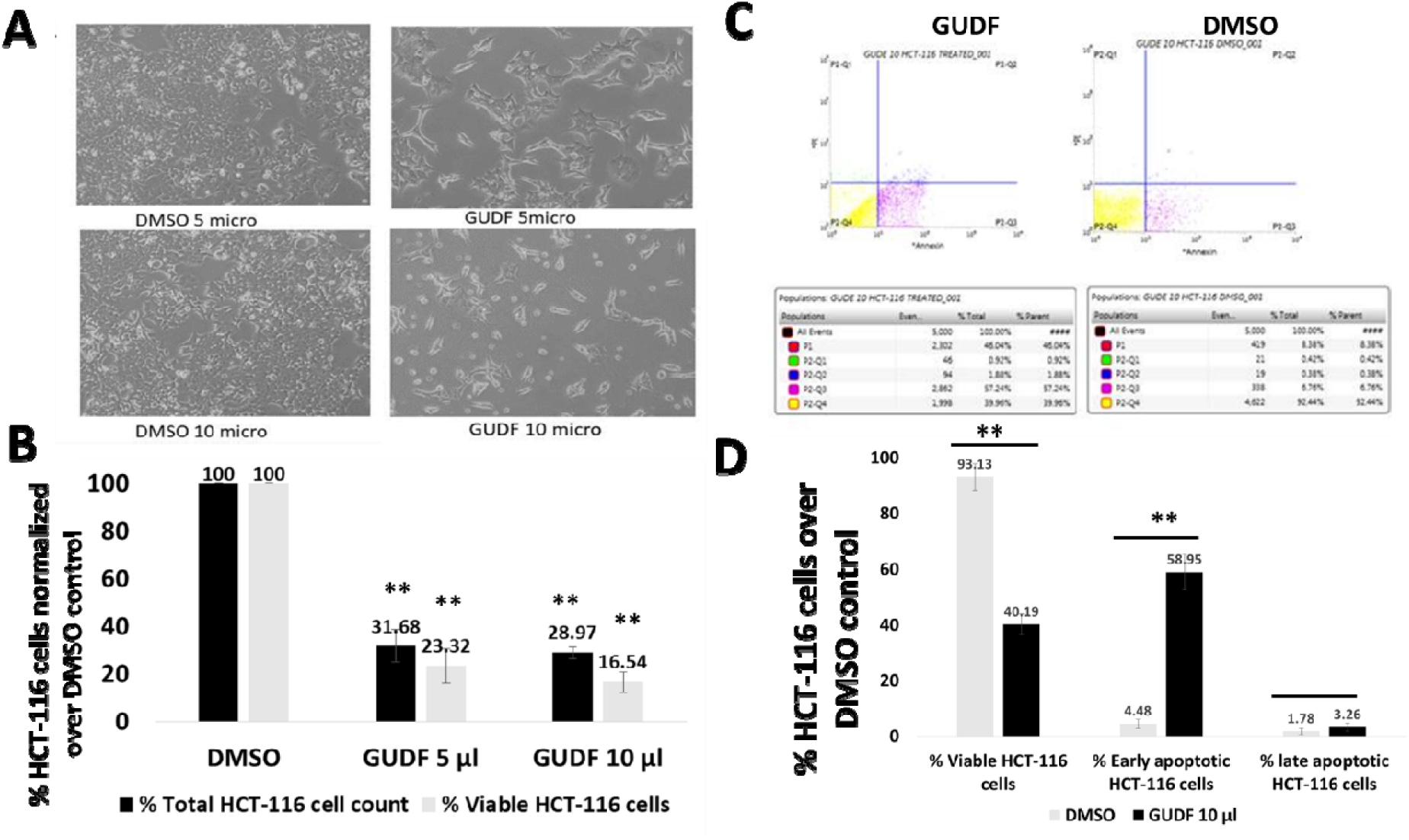
GUDF enriched with tripeptides show a clear retardation of proliferation of HCT-116 cells and significant presence of early apoptotic cell death. But, complete apoptotic cell death of HCT-116 was not observed. HCT-116 cells were treated with GUDF and solvent control DMSO for 72 hr. Then, harvested cells were subjected to Trypan blue dye exclusion assay to estimate total cell and viable cells. (A) A representative phase contrast image of HCT-116 cells treated with DMSO and GUDF i imaged at 100 X magnification. (B) This bar graph shows the percentage reduction of total HCT-116 cells and viable HCT-116 in case of GUDF treated cells normalized to DMSO control. Estimation of loss of viability and presence of apoptotic cell death in HCT-116 cells i determined by PI/annexin V staining assay. At the end of treatment as mentioned above, harvested HCT-116 cells were subjected to PI/annexin V staining and analyzed by flow cytometer. Data represent the cells stained with PI and Annexin V conjugated with FITC for the analysis of apoptotic cells in HCT-116 cells. (C) A representative PI/Annexin stained scatter plot of HCT-116 cells treated with DMSO and GUDF DOX are shown. (D). A bar graph represents % of total viable HCT-116 cells and % of early and late apoptotic HCT-116 cells. Data are represented as mean ± SD. Each experiment was conducted independently three times. The bar graph without an asterisk denotes that there is no any significant difference compared to DMSO control. * Significantly different from DMSO control at the P-value < 0.05. ** Significantly different from DMSO control at P-value < 0.01.

## IDENTIFICATIONS OF INTRACELLULAR TRIPEPTIDES IN HCT-116 cells

Based on the above observations, we proposed to conduct the intracellular metabolite profiling of HCT-116 cells treated with GUDF enriched with tripeptides and DMSO control. We hoped that information on the components of GUDF enriched with tripeptides that have entered the intracellular compartment of HCT-116 may pave the way for the next set of experiments to reveal detailed understanding of the cause behind the proliferation arrest. In this direction, we employed a novel methodology and process to study the intracellular metabolite profiling of HCT-116 cells that included the purification of hypotonically prepared cell lysates by using the vertical tube gel electrophoresis (VTGE) system (45,46,53).

The analysis of data based on LC-HRMS of purified intracellular metabolites of HCT-116 cells treated by GUDF enriched with tripeptides and DMSO suggested the distinct set of tripeptides. The major abundant tripeptides in DMSO treated HCT-116 cells are found as Ser-Trp-Lys, Glu-Glu-Gln, Glu-Glu-Lys and Ser-Leu-Ser. The details of these tripeptides along with characteristics product ion spectra is given in Table 1A and Figure S4, S5, S6 and S8). On the other side, GUDF enriched with tripeptides treated HCT-116 cells revealed another set of tripeptides as Glu-Glu-Arg, Gly-Arg-Pro, Gln-Lys-Arg, Glu-Glu-Lys, Trp-Trp-Val and their detailed mass ion spectra and properties are provided in Table 1B and Figure 3, Figure S2, S3, S6 and S7.

**Table 1A.** List of DMSO treated HCT-116 cells intracellular tripeptides analyzed by LC-HRMS and their molecular details.

| Sr.<br>No | Name of<br>Tripeptides | Formula | +ESI EIC | m/z | Mass | Polarity | CHEBI-ACCESSION<br>ID |
| --- | --- | --- | --- | --- | --- | --- | --- |
| 1 | Ser-Trp-Lys | C<sub>20</sub>H<sub>29</sub>N<sub>5</sub>O<sub>5</sub> | 402.2080<br>505.3092<br>293.1883 | 402.208 | 419.2115 | Positive | CHEBI:144474 |
| 2 | Glu-Glu-Gln | C<sub>15</sub>H<sub>24</sub>N<sub>4</sub>O<sub>9</sub> | 449.1525<br>450.1555<br>451.1575 | 449.1514 | 404.1532 | Positive | CHEBI-144559 |
| 3 | Glu-Glu-Lys | C<sub>16</sub>H<sub>28</sub>N<sub>4</sub>O<sub>8</sub> | 387.1874<br>304.2964<br>467.1548 | 387.1885 | 404.1919 | Positive | CHEBI:144461 |
| 4 | Ser-Leu-Ser | C<sub>12</sub>H<sub>23</sub>N<sub>3</sub>O<sub>6</sub> | 199.0685<br>293.0968<br>480.1727 | 328.1503 | 305.1588 | Positive | CHEBI-144475 |

**Table 1B.**
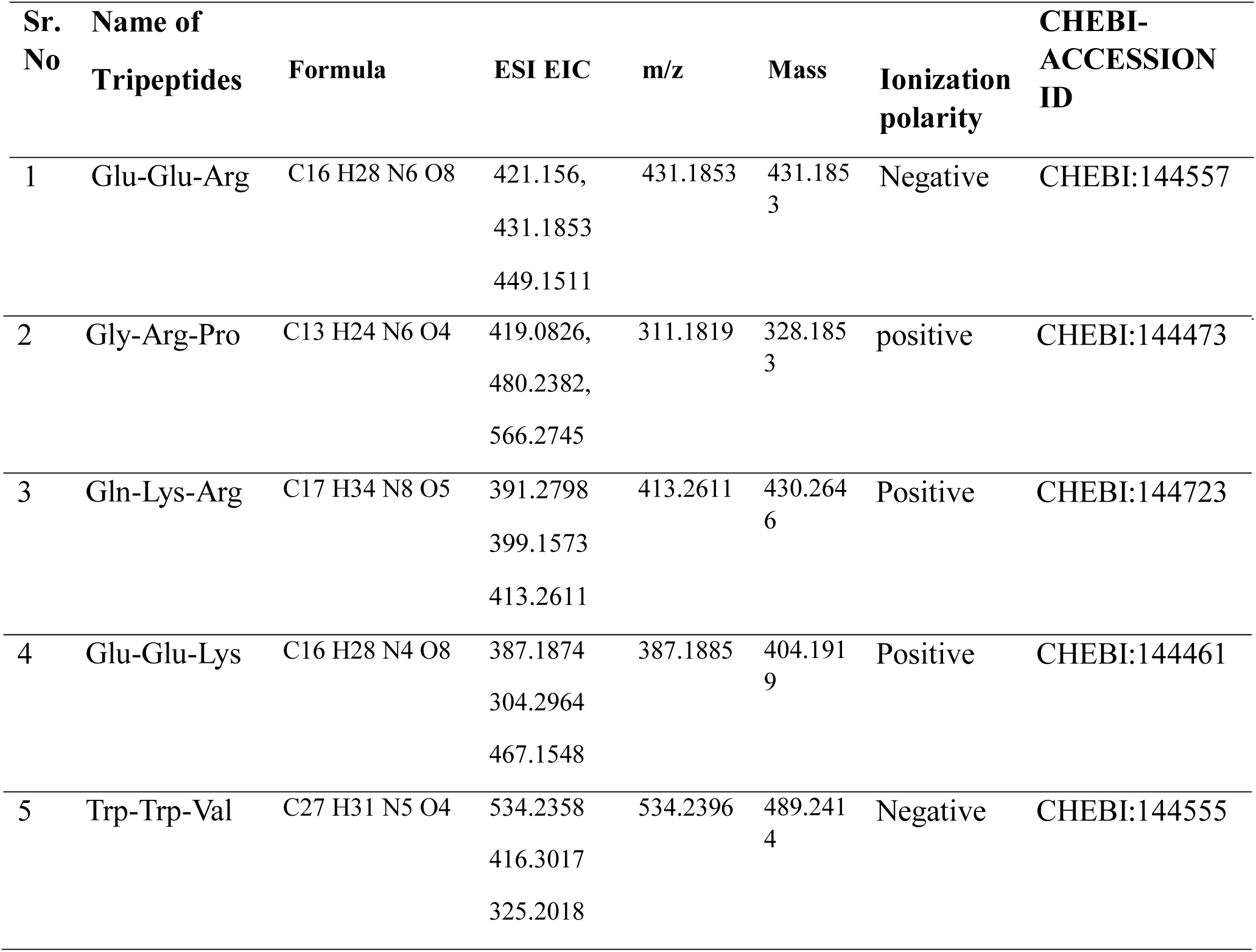
List of intracellular tripeptides in HCT-116 cells treated by GUDF enriched with tripeptides and these metabolites were analyzed by LC-HRMS with details of mass and +ESI EIC and - ESI EIC.

| Sr.<br>No | Name of<br>Tripeptides | Formula | ESI EIC | m/z | Mass | Ionization<br>polarity | CHEBI-ACCESSION<br>ID |
| --- | --- | --- | --- | --- | --- | --- | --- |
| 1 | Glu-Glu-Arg | C<sub>16</sub>H<sub>28</sub>N<sub>6</sub>O<sub>8</sub> | 421.156,<br>431.1853<br>449.1511 | 431.1853 | 431.1853 | Negative | CHEBI:144557 |
| 2 | Gly-Arg-Pro | C13 H24 N6 O4 | 419.0826,<br>480.2382,<br>566.2745 | 311.1819 | 328.185<br>3 | positive | CHEBI:144473 |
| 3 | Gln-Lys-Arg | C17 H34 N8 O5 | 391.2798<br>399.1573<br>413.2611 | 413.2611 | 430.264<br>6 | Positive | CHEBI:144723 |
| 4 | Glu-Glu-Lys | C16 H28 N4 O8 | 387.1874<br>304.2964<br>467.1548 | 387.1885 | 404.191<br>9 | Positive | CHEBI:144461 |
| 5 | Trp-Trp-Val | C27 H31 N5 O4 | 534.2358<br>416.3017<br>325.2018 | 534.2396 | 489.241<br>4 | Negative | CHEBI:144555 |

**Figure 3:**
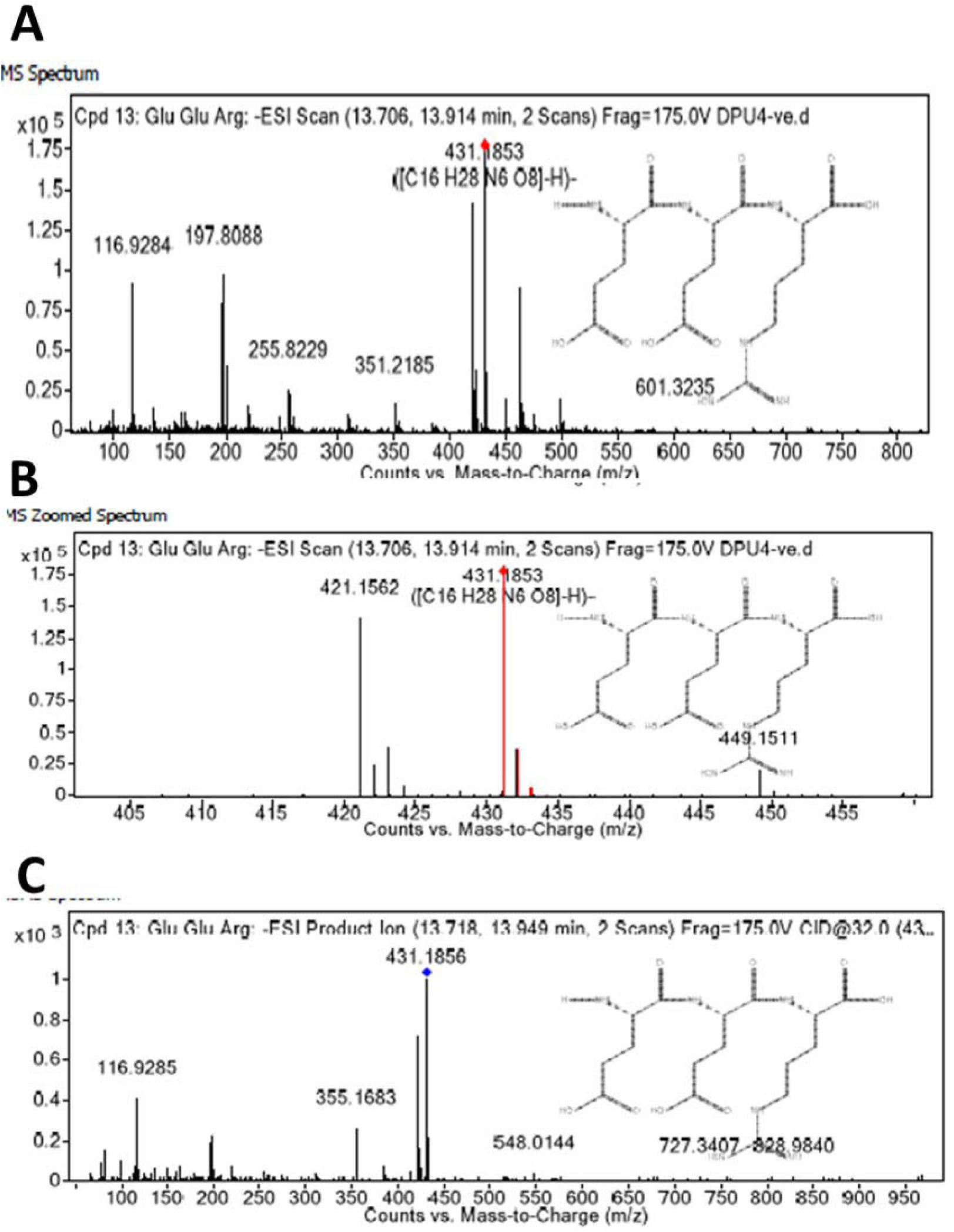
LC-HRMS analysis of intracellular metabolite analysis of HCT-116 cells treated with GUDF reveals abundance of Glu-Glu-Arg and other tripeptides. Hypotonically prepared cell lysate of HCT-116 cells treated with GUDF was purified by employing a novel methodology VTGE and analyzed by LC-HRMS technique in negative ESI-MS mode. (A) MS spectrum of total negative ESI scan. (B). MS spectrum of zoomed negative ESI scan. (C). MS/MS negative ESI product ion spectra.

The abundance of these tripeptides in the intracellular compartment of treated HCT-116 cells raised questions on the biological relevance of these tripeptides. In literature, data suggest that different types of tripeptides based on their amino acids residues may contribute either as pro-proliferation or anti-proliferation (29-37). Among the reported tripeptides, the majority of tripeptides are attributed with basic amino acids such as Arg, Lys, His. In this study, the nature of intracellular tripeptides in HCT-116 cells treated GUDG enriched with tripeptides are mostly basic in nature that contained Arg, Gln and Gly. Serum components of culture are known to contain selected tripeptides that contribute towards cellular growth and proliferation. An interesting old paper reported on the ability of a tripeptide Gly-His-Lys, a human plasma constituent to chelate with metal ions during growth conditions and may modulate the growth potential of cancer cells (30). In summary, during normal growth conditions, tripeptides components from media may enter in the intracellular compartment of HCT-116 cells. On the other hand, GUDF treated HCT-116 cells raises the possibility of stress condition within the cells and uptake of anti-growth and proliferation tripeptides might show the arrest of proliferation. Here, authors would like to mention that detection of unique set of intracellular metabolites in HCT-116 cells by the mentioned novel methodology is novel and first time. Therefore, abundance of distinct set of tripeptides in DMSO and GUDF treated HCT-116 cells suggests the role in intracellular signaling that support the growth and proliferation of cancer cells.

## MOLECULAR DOCKING OF TRIPEPTIDES WITH AP-1 TRANSCRIPTIONAL COMPLEX

Based on the cellular effects and nature of intracellular peptides in HCT-116 cells, we proceeded to evaluate the relevance of intracellular peptides with observed proliferation arrest. We looked into literature on the molecular basis of proliferation arrest and one strong mechanism is in the form of disruption of transcriptional complex in cancer cells. Here, it is important to mention that selected tripeptides and inhibitors are known to serve as a chemical inhibitor to block the transcriptional complex such as AP-1, c-Myc and FOXO3 transcription factors (14-25).

Particularly, AP-1 involves heterodimer structure of c-Jun and c-Fos that clamps on consensus sequence (3,10-13). Furthermore, c-Jun was initially identified as a novel oncoprotein of avian sarcoma virus and c-Fos showed homology with v-fos oncogenes that is known to induce osteosarcoma. Interestingly, AP-1 binding site was discovered as 12-O-Tetradecanoylphorbol-13-acetate (TPA) response element that possess consensus heptamer sequence 5’-TGAGTCA-3’ as binding pocket for heterodimer c-Jun and c-Fos (3,10-13). In view of disruption of AP-1 transcriptional complex, heptamer sequence 5’-TGAGTCA-3’ within TPA of AP-1 complex is of great importance, since this is the coding strand which is responsible for transcription of a set of genes involved proliferation and differentiation.

Currently, AutoDock Vina is employed for molecular docking experiments with reference to notable attributes such as better accuracy that predict binding patterns, shortened run time and efficient search of potential energy surfaces and higher reproducibility (26). In this paper, target AP-1 transcriptional complex was retrieved from the PDB ID (1FOS) and detailed crystal structure show the clamping of heterodimer c-Jun:c-Fos on 20 nucleotide TPA response element including heptamer AP-1 consensus site (5′-TGAGTCA-3′) (10).

In this way, AutoDock Vina was employed to evaluate selected DMSO treated HCT-116 cells intracellular tripeptides (Ser-Trp-Lys, Glu-Glu-Gln, Glu-Glu-Lys, Ser-Leu-Ser) and GUDF treated HCT-116 cells (Glu-Glu-Arg, Gly-Arg-Pro, Gln-Lys-Arg, Glu-Glu-Lys, Trp-Trp-Val) against AP-1 transcriptional complex (PDB ID:1FOS). In the same design of molecular docking study, authors assessed the molecular binding patterns of known inhibitors (NDGA, curcumin), intercalating agent (ethidium bromide) and DNA binding dye (Hoechst 33342).

An interesting tripeptide Glu-Glu-Arg detected in GUDG treated HCT-116 cells showed highest binding affinity at −9.1 Kcal/Mol among tested tripeptides and known inhibitors (Figure 4A and 4B). It is important to note that along with high docking affinity of Glu-Glu-Arg, this peptide interact within the heptamer AP-1 consensus site (5′-TGAGTCA-3′) of AP-1 transcriptional complex by binding to DG8, DA9, DA31, DG30 and c-Fos (Ser-154) chain and c-Jun chain (Arg-279) (Figure 4C and 4D). The ability of Glu-Glu-Arg to achieve a strong and precise association with AP-1 transcriptional complex is reflected in the focused PyMol view with a total of thirteen polar bonds (Figure 4D, Table 4). Another tripeptide Gly-Arg-Pro possess specific docking ability within the AP-1 transcriptional complex with a binding energy at −8.8 Kcal/Mol (Figure 5A and 5B). This tripeptide makes access within the heptamer AP-1 consensus site (5′-TGAGTCA-3′) with specific binding to DG8, DG30, c-Fos (Ser-154, Arg-155) and c-Jun (Ala-275) (Figure 5C and 5D). Besides these two arginine containing tripeptides, data suggest the specific binding ability of Gln-Lys-Arg by showing molecular interaction with A-chain (DA9, DG10, DT11, DC12, DA13) and B-chain (DC32, DT33, DA35) of heptamer consensus sequence (5′-TGAGTCA-3′) of AP-1 complex (Figure 6A, 6B, 6C and 6D).

**Figure 4.**
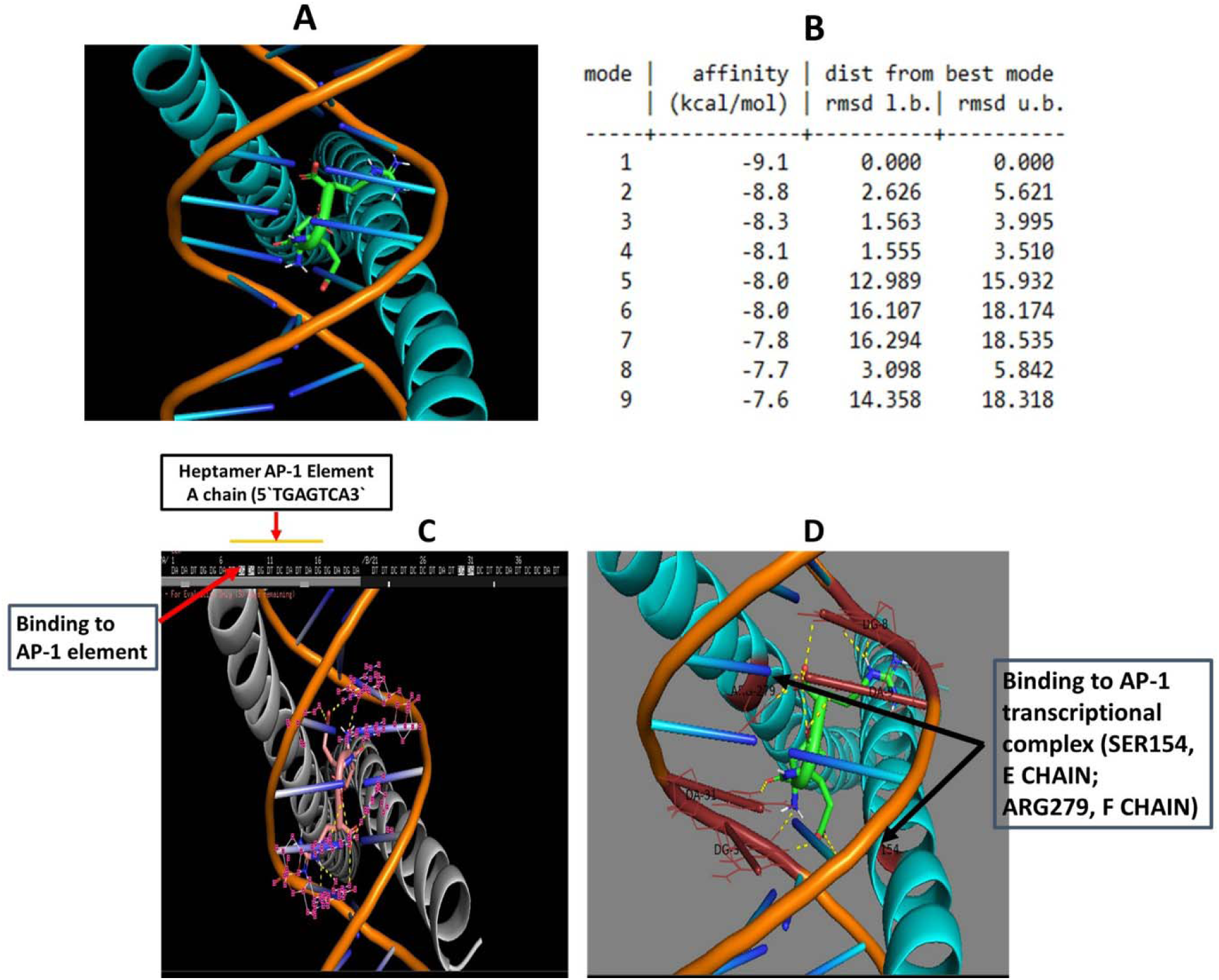
GUDF tripeptide Glu-Glu-Arg shows a strong binding to AP-1 consensus site (5′-TGAGCTCA-3′, also known as TPA-responsive element and ternary AP-1 transcriptional complex c-Jun:c-Fos:DNA. (A) A computer generated PyMol view of Glu-Glu-Arg ligand bound to ternary complex of c-Jun-c-Fos:DNA. (B) Molecular docking energy estimated by AutoDock Vina during the interaction of Glu-Glu-Arg with c-Jun-c-Fos:DNA ternary complex with different rmsd l.b. and u.b. value. (C) Emphasized view by PyMol showing the specific binding of Glu-Glu-Arg within the heptamer AP-1 consensus site (5′-TGAGTCA-3′ of AP-1 transcriptional complex (DG8, DA9, DA31, DG30). (D) In depth image of PyMol view that display the binding of Glu-Glu-Arg to E-chain (c-Fos) amino acid residue Ser-154 and F-chain (c-Jun) amino acid residue Arg-279.

**Figure 5.**
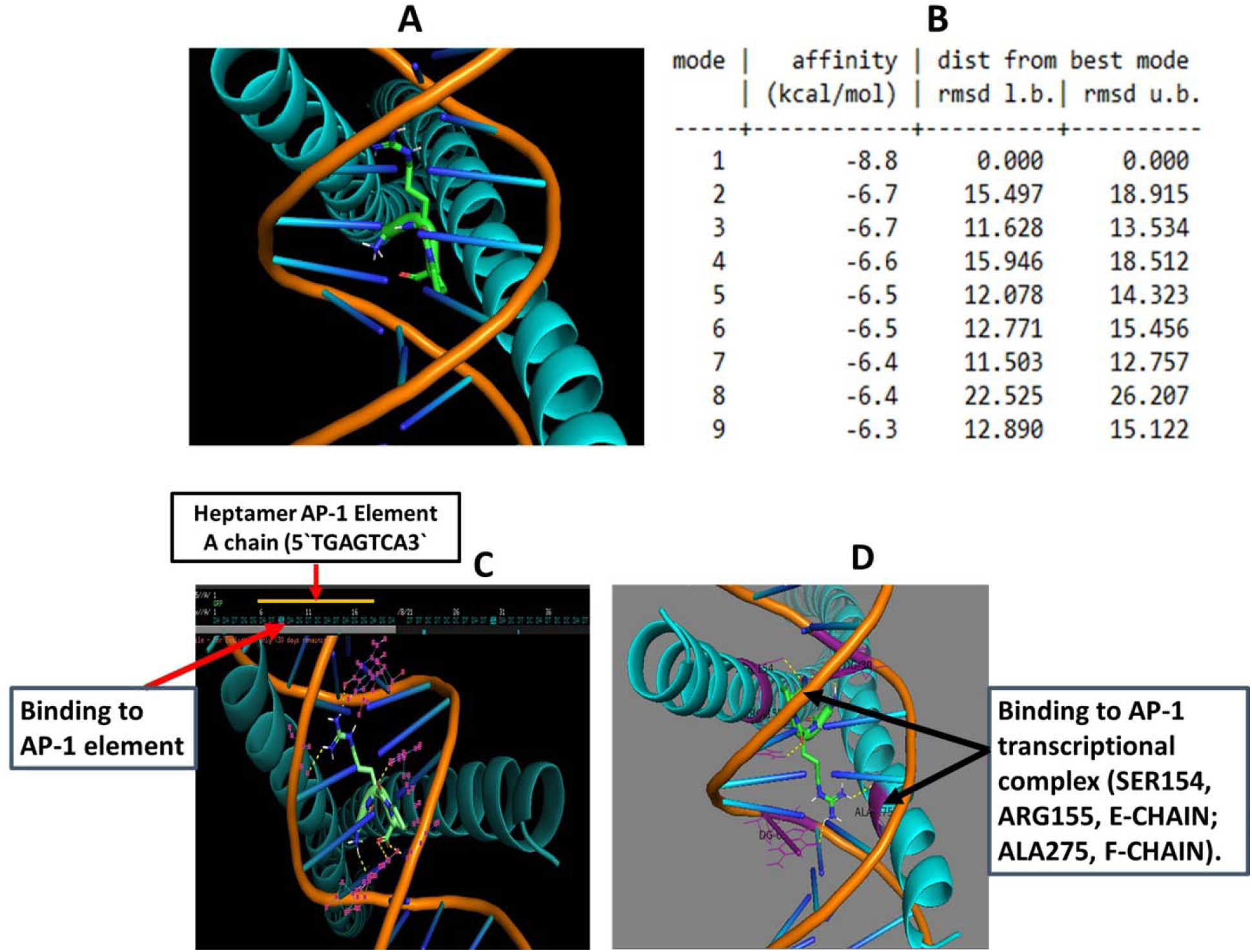
GUDF derived tripeptide Gly-Arg-Pro demonstrates strong molecular interaction within the AP-1 transcriptional complex with a docking energy value at −8.8 Kcal/Mol. Figure 4. GUDF tripeptide Gly-Arg-Pro displays a specific and clear molecular interaction to AP-1 consensus site (5′-TGAGCTCA-3′) within the ternary transcriptional complex of c-Jun:c-Fos:DNA. (A) A computer generated PyMol view of Gly-Arg-Pro ligand bound to ternary complex of c-Jun-c-Fos:DNA. (B) Molecular docking energy estimated by AutoDock Vina during the interaction of Gly-Arg-Pro with c-Jun-c-Fos:DNA ternary complex with different rmsd l.b. and u.b. value. (C) Emphasized view by PyMol showing the specific binding of Gly-Arg-Pro within the heptamer AP-1 consensus site (5′-TGAGTCA-3′) of AP-1 transcriptional complex (DG8, DG30). (D) In depth image of PyMol view that display the binding of Gly-Arg-Pro to E-chain (c-Fos) amino acid residue Ser-154, Arg-155 and F-chain (c-Jun) amino acid residue Ala-275.

**Figure 6.**
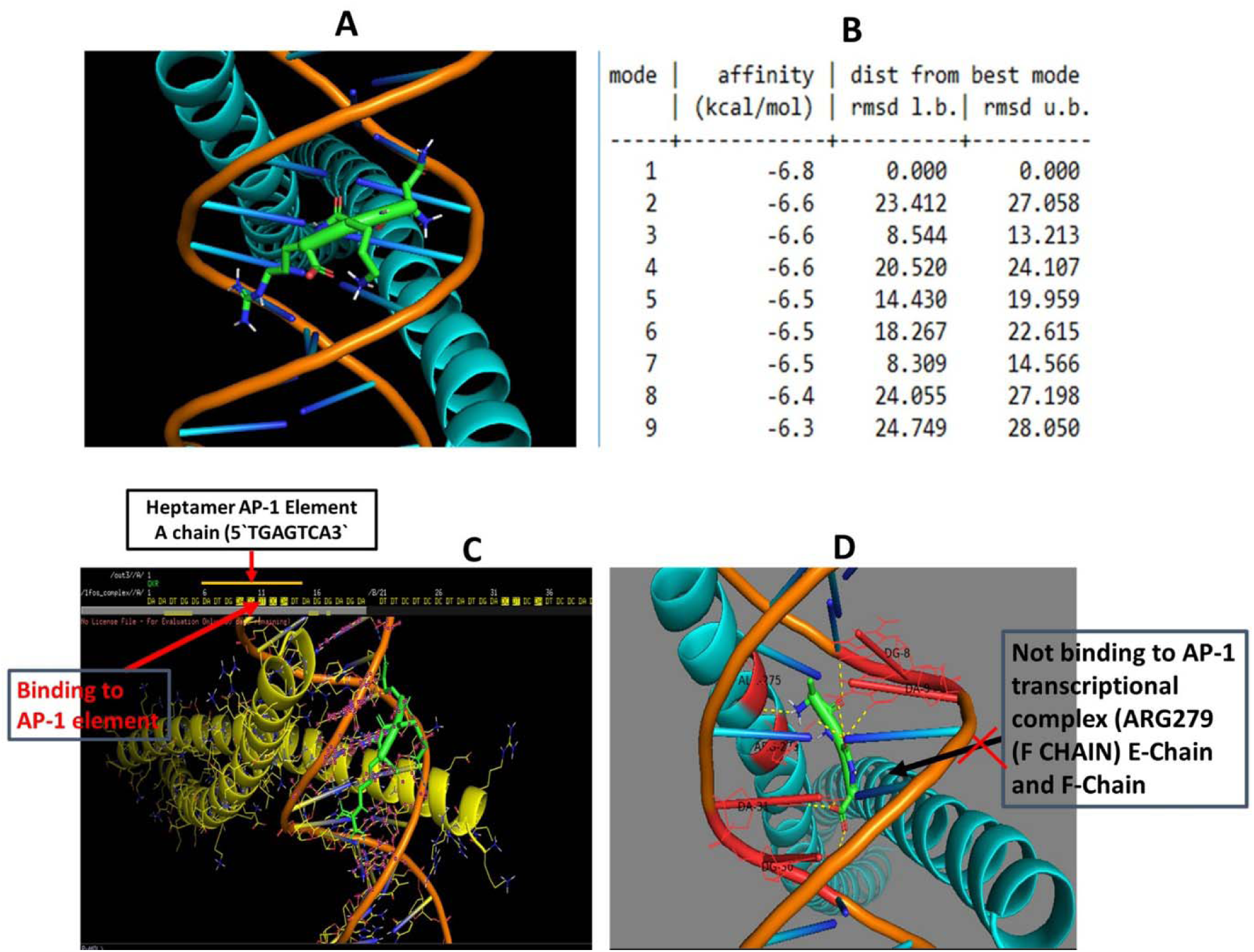
GUDF derived tripeptide Gln-Lys-Arg displays specific interaction within the AP-1 transcriptional complex with a docking energy value at −6.8 Kcal/Mol with a maximum number of polar bonds. (A) A computer generated model by PyMol displays the binding between tripeptide Gln-Lys-Arg, and AP-1 transcriptional complex. (B). Molecular docking energy data obtained by AutoDock Vina based molecular interaction between Gln-Lys-Arg and c-Jun-c-Fos:DNA ternary complex show the docking affinity value with reference to distance from rmsd u.b. and best mode rmsd l.b. with a maximum value at 6.8 Kcal/Mol. (C) Emphasized image generated by PyMol depicts the clear and specific binding to heptamer AP-1 consensus site (5′-TGAGTCA-3′ of AP-1 transcriptional complex (DA9, DG10, DT11, DC12, DA13, DC32, DT33, DA35). (D). Additional in depth pose generated by PyMol supports the non-association of Gln-Lys-Arg with c-Jun and c-Fos polypeptide heterodimer of AP-1 complex.

Data clearly indicate the among all tested tripeptides, Glu-Glu-Arg, Gly-Arg-Pro and Gln-Lys-Arg are highly efficient in binding to c-Fos:c-Jun:DNA complex with a specificity within the heptamer AP-1 consensus site (5′-TGAGTCA-3′). In a comparative analysis with known AP-1 complex inhibitor (NDGA, curcumin), and DNA binding agent (ethidium bromide and Hoechst 33342), these novel tripeptides including Glu-Glu-Arg, Gly-Arg-Pro and Gln-Lys-Arg demonstrates better abilities in terms of docking energy, specificity within the AP-1 consensus site (5′-TGAGTCA-3′) and total number of polar bonds that destabilizes the AP-1 complex.

Besides Glu-Glu-Arg, Gly-Arg-Pro and Gln-Lys-Arg tripeptides, tripeptides such as Ser-Leu-Ser (−6.0 Kcal/Mol), Ser-Trp-Lys (−7.9 Kcal/Mol), Glu-Glu-Gln (−7.9 Kcal/Mol), Glu-Glu-Lys (−6.4 Kcal/Mol and Trp-Trp-Val (−8.3 Kcal/Mol) did not displayed appreciable specific binding to the heptamer AP-1 consensus site (5′-TGAGTCA-3′) of AP-1 transcriptional complex (Figure 8, Figure S9, Figure S10, Figure S11 and Figure S12, respectively). In addition to tripeptides, we have studied natural inhibitors such as NDGA and curcumin against AP-1 complex. Actually, NDGA is reported as an inhibitor of c-Fos:c-Jun:DNA complex and demonstrated anticancer effects. However, molecular docking study on NDGA over c-Fos:c-Jun:DNA complex is not available. Our molecular docking data substantiate the previous in vitro experimental evidence on the ability of NDGA that disrupt the AP-1 transcriptional complex (Table 2 and 3, Figure 7). This NDGA inhibitor showed strong binding with DG8 and DA9 within the heptamer AP-1 consensus site (5′-TGAGTCA-3′) of AP-1 transcriptional complex. NDGA is shown to bind to the Arg-279 that is at close proximity to Ser-278 and both amino acid residues are necessary for the clamping by c-Jun upon TPA response element. Detailed analysis of molecular docking by NDGA reveals a binding energy at −7.6 Kcal/Mol and three polar bonds are observed. Another natural anticancer and inflammatory compound, curcumin demonstrates binding affinity to the DNA structure with binding energy value at −7.8 Kcal/Mol. Interestingly, PyMol based emphasize view of docking complex between curcumin and AP-1 complex, there is no specific destabilization of heptamer AP-1 consensus site (5′-TGAGTCA-3′) of AP-1 transcriptional (Table 2, Table 3 and Figure S13).

**Table 2.**
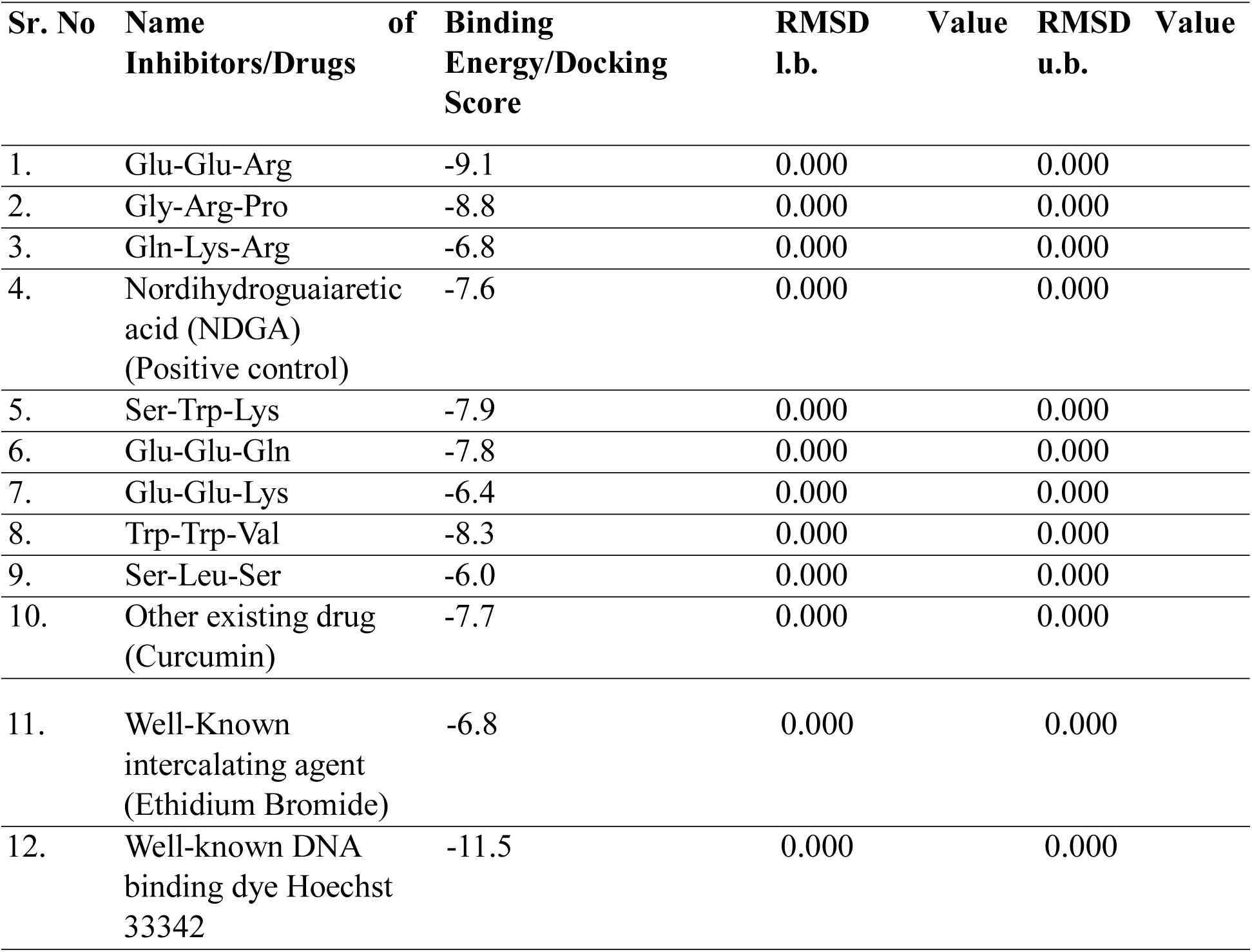
List of inhibitors/drugs evaluated for inhibition of AP-1 transcriptional complex with their binding energy score. Here, molecular interaction studies between selected tripeptides/inhibitor and c-Jun:c-Fos:DNA (PDB ID:1FOS) was studied by using AutoDock Vina.

**Table 3.**
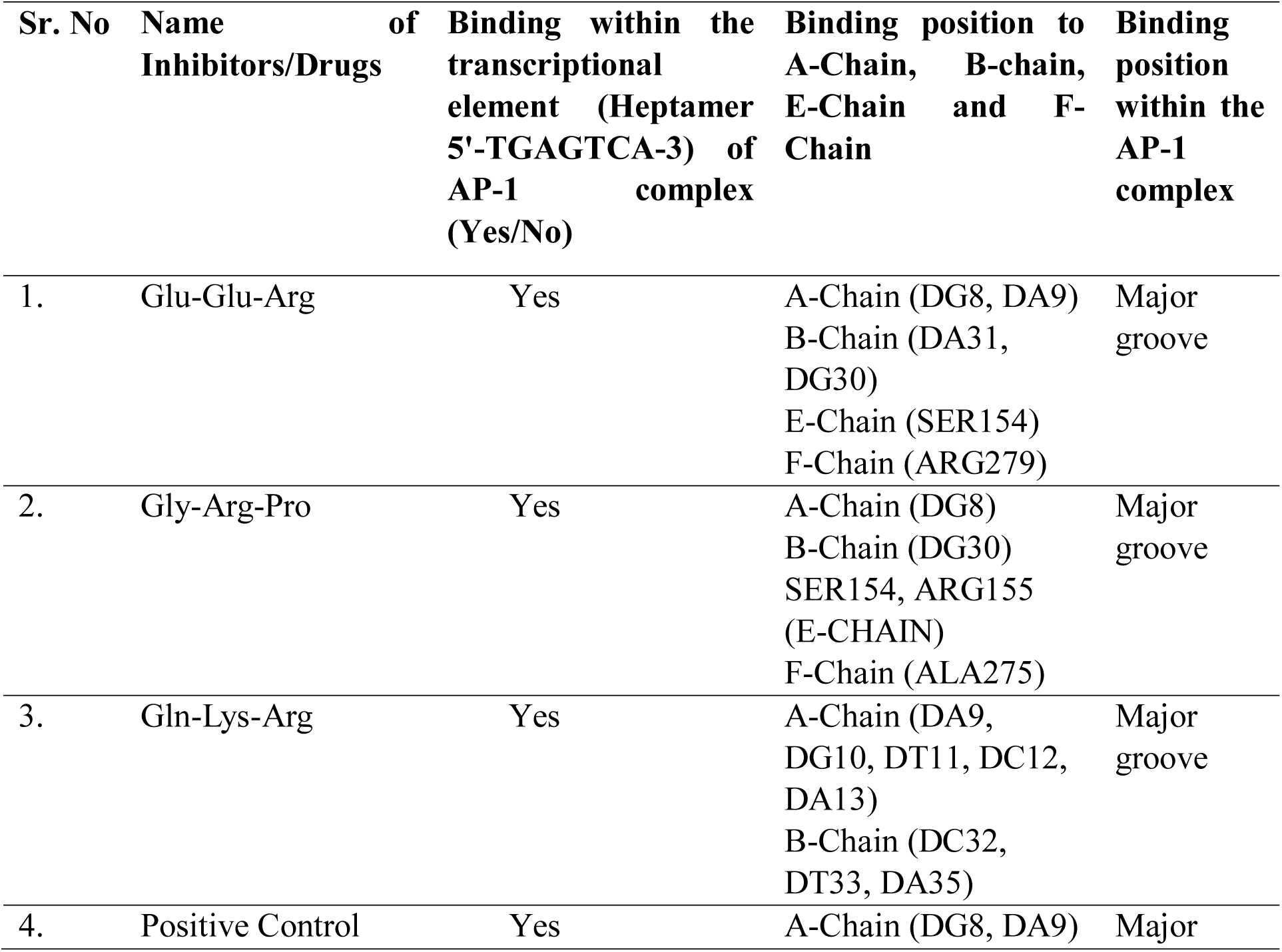
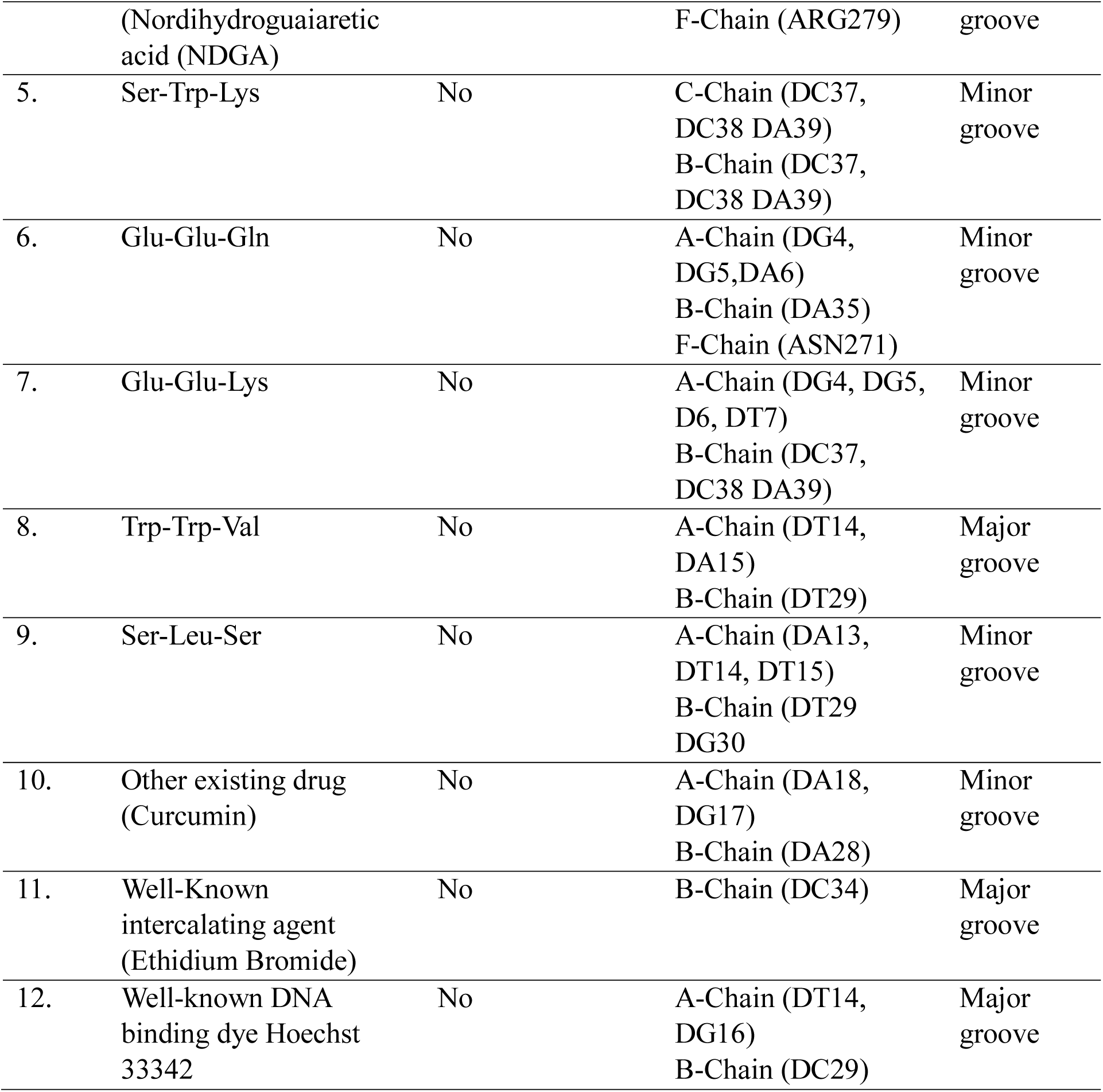
List of tripeptides/inhibitors/drugs that bind within the heptamer 5’-TGAGTCA-3’ element within the AP-1 transcriptional complex with their binding properties. Here, PyMol based view of molecular interaction model between selected tripeptides/inhibitor and c-Jun:c-Fos:DNA (PDB ID:1FOS) was studied.

**Figure 7.**
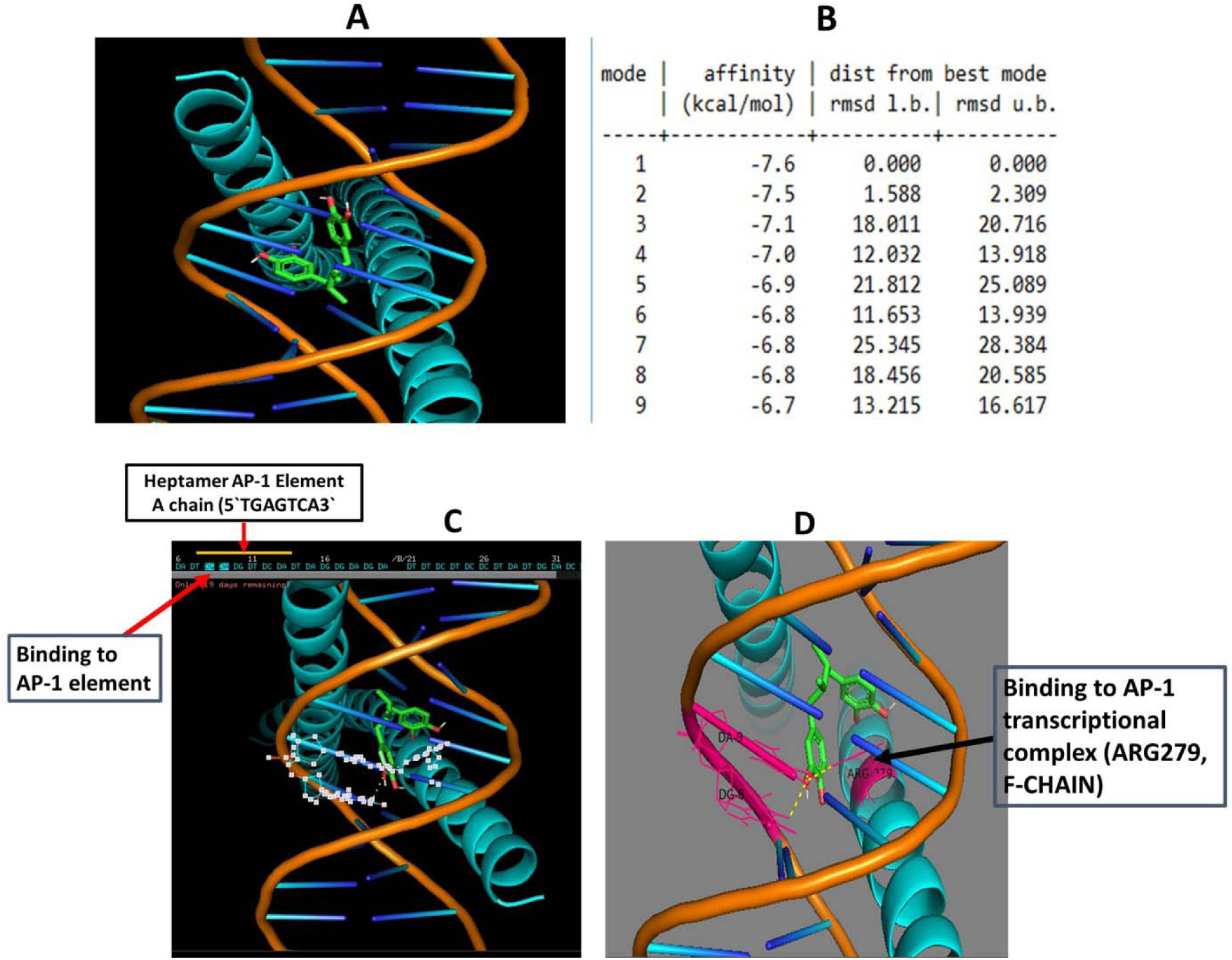
A known positive control Nordihydroguaiaretic acid (NDGA) shows binding to AP-1 transcriptional complex with a docking energy value at −7.6 Kcal/Mol. (A) A computer generated model by PyMol displays the binding between NDGA, a known inhibitor of AP-1 transcriptional complex. (B). Data generated from AutoDock Vina based molecular interaction between NDGA and c-Jun-c-Fos:DNA ternary complex show the docking affinity value with reference to distance from rmsd u.b. and best mode rmsd l.b. with a maximum value at 7.6 Kcal/Mol. (C) Emphasized image generated by PyMol depicts the clear and specific binding to heptamer AP-1 consensus site (5′-TGAGTCA-3′) of AP-1 transcriptional complex (A-chain). (D). Additional in depth pose generated by PyMol supports the bidding to the F-chain (c-Jun) polypeptide at Arg-279 amino acid residue.

**Figure 8.**
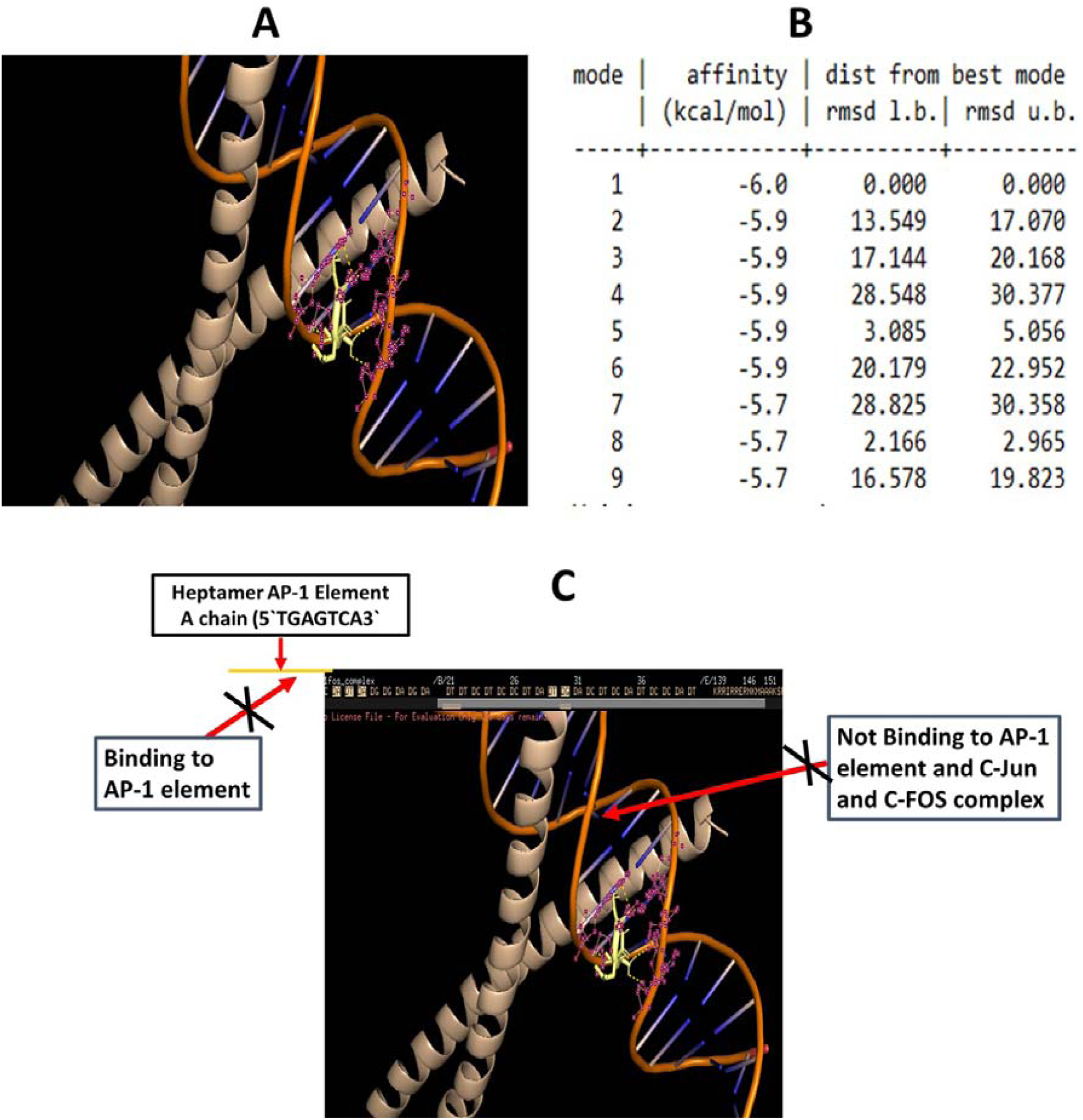
An intracellular tripeptide Ser-Leu-Ser from DMSO treated HCT-116 cells does not show a specific binding to AP-1 transcriptional complex and low docking energy value is at −6.0 Kcal/Mol. (A). A PyMol view of docking between tripeptide Ser-Leu-Ser and c-Jun:c-Fos:DNA ternary complex. (B) Value on docking affinity obtained during AutoDock Vina based molecular interaction between tripeptide Ser-Leu-Ser and c-Jun:c-Fos:DNA ternary complex. (C). In depth image of PyMol generated model shows that tripeptide Ser-Leu-Ser is not able to bind within the heptamer AP-1 consensus site (5′-TGAGTCA-3′) of AP-1 transcriptional complex. Also, tripeptide Ser-Leu-Ser does not bind to E-chain (c-Fos) and F-chain (c-Jun) heterodimer complex within the AP-1 transcriptional complex.

In addition to NDGA and curcumin, ethidium bromide, a known DNA intercalating agent displays non-specific binding to DNA with a binding affinity of −7.6 Kcal/Mol, other than heptamer AP-1 consensus site (5′-TGAGTCA-3′) of AP-1 transcriptional complex (Table 2, Table 3 and Figure S14). Another known DNA binding dye Hoechst 33342 also shows non-specific binding to DNA as expected with a strong binding affinity at −11.5 Kcal/Mol and this dye is not able to destabilize the AP-1 (Table 2, Table 3 and Figure S15).

An argument may be raised to understand the relevance of testing potential tripeptides against the AP-1 transcriptional complex. Actually, the biological relevance of dipeptide and tripeptide is suggested in view of basic understanding on protein-protein and protein DNA interactions. In cellular landscape, protein-protein and protein-DNA interaction hold keys for various cellular events such DNA replication, transcription and translation process (3-12). Therefore, limited approaches are noticed to unravel the biological propensity of conserved tripeptide motifs towards DNA substrates that may be specific or non-specific binding.

In essence, the binding of tripeptides with different chemistry are seen as inducers of DNA damage and also as non-covalent binding to the transcriptional response element as an inhibitor (18-25). In this paper, we examined the binding capabilities of selected tripeptides and that is based on the basic understanding that mostly di-peptide and tripeptides are crucial regions on DNA binding proteins including key transcription factors such as AP-1 and MYC (3-6,9-12). Furthermore, limited findings have attempted to look for chemical approaches to disrupt the binding between transcription factors and their target response binding element mostly upstream of set genes being controlled by transcriptional processes. Among potential chemical inhibitors, NDGA, curcumin, Arg-Pro-Arg, N-(3-acetamidophenyl)-2-[5-(1H-benzimidazol-2-yl)pyridin-2-yl]sulfanylacetamide and T-5224 are reported for their inhibitory potential on transcriptional complex (19-25). Besides these chemicals, natural compounds such as dihydroguaiaretic acid that was initially extracted from the aryls of Myristica fragrans, nordihydroguaiaretic acid (NDGA) and curcumin are shown to disrupt heterodimer of c-Jun:c-Fos that bind to AP-1 response element (14-18).

These chemicals as disruptors of AP-1 transcriptional complexes are studied by designing in-silico, in vitro and in vivo experimental approaches. In this paper, molecular docking data on known chemicals such as NDGA and curcumin are presented and that precisely show the binding of NDGA is within the heptamer AP-1 consensus site (5′-TGAGTCA-3′) at DG8, DA9 of AP-1 transcriptional complex. Besides these chemicals as an inhibitor of AP-1 transcriptional complex, molecular docking data is lacking on the novel tripeptides that show the specific and strong binding to heptamer AP-1 consensus site (5′-TGAGTCA-3′) of AP-1 transcriptional complex.

In a proof of concept, our selected tripeptides such as Glu-Glu-Arg (DG8, DA9, DA31, DG30), Gly-Arg-Pro (DG8, DG30) and Gln-Lys-Arg (DA9, DG10, DT11, DC12, DA13, DC32, DT33, DA35) binds specifically to AP-1 complex. Based on the above similar nature of binding to consensus sequence of AP-1 complex, these novel set of tripeptides Glu-Glu-Arg, Gly-Arg-Pro and Gln-Lys-Arg are suggested as an option for chemical inhibition of AP-1 transcriptional complex and a potential candidate for antiproliferative agents.

In our findings, Glu-Glu-Arg tripeptides show strong and precise binding to the specific amino acid residue Ser-154 (c-Fos) and Arg-279 (c-Jun). Additionally, Gly-Arg-Pro precisely interacts with Ser-154, Arg-155 (c-Fos) and Ala-275 (c-Jun) amino acid residues. The biological relevance of these observations is linked with the basic structure of the AP-1 transcriptional complex. In essence, the bZIP domain of c-Jun and c-Fos that together create a heterodimer is known to bind to a 20 nucleotide DNA duplex that encompass heptamer consensus sequence (5′-TGAGTCA-3′) (3,10-13). The detailed studies on the c-Jun:c-Fos:DNA ternary complex revealed that basic regions of heterodimer bind to the base pairs of AP-1 consensus site. The residues in this ternary complex are known to contribute to high affinity of binding and desired interactions of c-Jun:c-Fos:DNA that achieve the stable transcriptional complex for gene expression (3,10-13). Based on the crystallographic and mutation studies, basic amino acid residues of c-Fos including Arg-146, Asp-147, Ala-150, Ala151, Lys-153, Ser-154, Arg-155 and Arg-268, Asp-271, Arg-272, Ala-274, Ala-275, Lys-277, Ser-278, Arg-279 of c-Jun binds to AP-1 DNA sequence in an asymmetrically manner and form a stable ternary transcriptional complex (10,11). Therefore, key amino acid residues including Ser-154, Arg-155 (c-Fos) and Ala-275, Arg-279 (c-Jun) are bound by the tripeptides in this peptides and these amino acid residues are crucial for the stable and efficient AP-1 transcriptional complex. Additionally, a known inhibitor of AP-1 transcriptional complex NDGA also binds to the c-Fos (Ser-154) chain and c-Jun chain (Arg-279) similar to claimed tripeptides in this study.

Based on the comparative binding affinity of selected tripeptides, known inhibitors, chemicals, DNA binding dye, Glu-Glu-Arg, Gly-Arg-Pro and Gln-Lys-Arg show better or equivalent binding affinity to the consensus sequence of AP-1 complex. Interestingly, details of polar bonds that allow these tripeptides (13, 6, 11) to destabilize the AP-1 complex are higher than the known inhibitor NDGA (3) in terms of number of polar bonds and precisely distance of polar bonds are equal in case of selected tripeptides (1.9-2.8 compared to NDGA (1.9-2.3) (Table 4). Furthermore, Glu-Glu-Arg, Gly-Arg-Pro and Gln-Lys-Arg tripeptides bind to the major groove of the heptamer AP-1 consensus site (5′-TGAGTCA-3′) of AP-1 transcriptional complex and this interaction matches with the major groove binding by NDGA, a known inhibitor of AP-1 complex (Table 3). Therefore, the claims made on these selected tripeptides that are based on the in silico approach are strengthened with respect to the pattern of molecular binding displayed by known inhibitors of AP-1 complex.

**Table 4.**
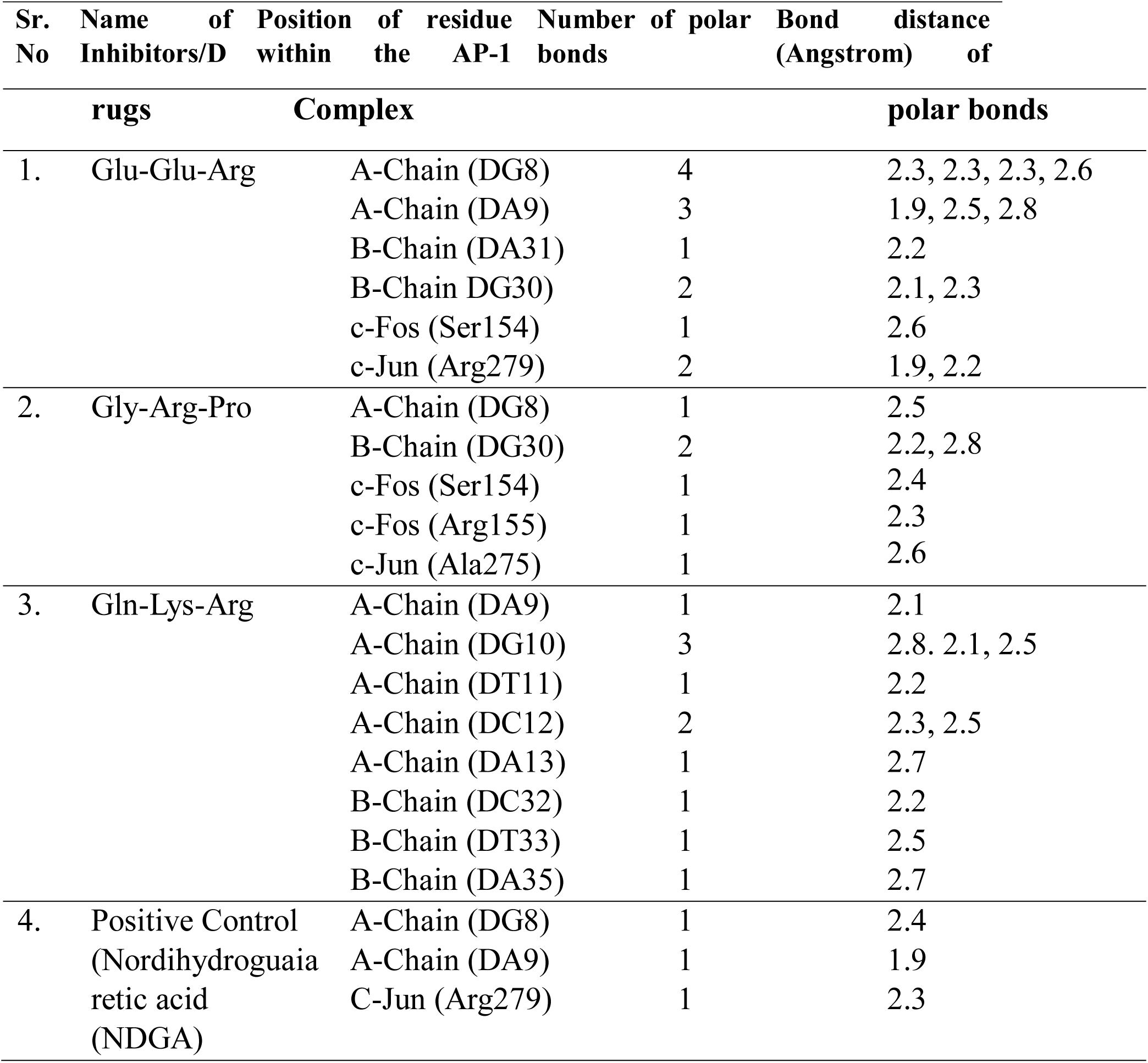
Details of polar bonds within the heptamer 5’-TGAGTCA-3’ consensus sequence AP-1 transcriptional complex with their number and bond distance. Here, PyMol based view of molecular interaction model between selected tripeptides/inhibitor and c-Jun:c-Fos:DNA (PDB ID:1FOS) was studied.

Since, we show that abundance of tripeptides such as Glu-Glu-Arg, Gly-Arg-Pro and Gln-Lys-Arg are abundant in the intracellular compartment of HCT-116 cells that also showed the growth and proliferation arrest. Furthermore, molecular docking data confirm on strong and specific binding by Glu-Glu-Arg, Gly-Arg-Pro and Gln-Lys-Arg to the heptamer AP-1 consensus site (5′-TGAGTCA-3′) of AP-1 transcriptional complex and heterodimer c-Jun and C-Fos. These observations suggest the possibilities of the antiproliferative effects of Glu-Glu-Arg, Gly-Arg-Pro and Gln-Lys-Arg may be mediated through the efficient binding to the AP-1 transcriptional complex. Because, existing views support that selected tripeptides may work as anticancer agents by modulating distinct cancer cell signaling pathways including cell cycle progression, apoptosis and mitochondrial dysfunction (28-37). Based on existing understanding, diverse nature of tripeptides due to their inherent chemistry are involved in distinctive cellular processes. Among various cellular processes, the role of AP-1 transcription factor is well evident in proliferation and invasiveness of various types of cancer (28-37). Simultaneously, findings converge to suggest that inhibition of AP-1 transcriptional complexes force cancer cells towards arrest of cell cycle progressions (28-37). Interestingly, limited small molecular inhibitors are reported that show the affinity to block AP-1 transcriptional complexes and as a consequence proliferation in cancer cells. However, data is scarce that has attempted to investigate the novel tripeptides from biological sources that display specific binding abilities to transcriptional complex such as AP-1 by using molecular docking studies and supported with in vitro cell based observations. Our data warrants the potential of set tripeptides Glu-Glu-Arg, Gly-Arg-Pro and Gln-Lys-Arg that can inhibit the AP-1 transcriptional complex and may be explored as antiproliferative agents at preclinical and clinical levels. A summary of the proposed mechanism of these novel tripeptides is outlined in Figure 9.

**Figure 9.**
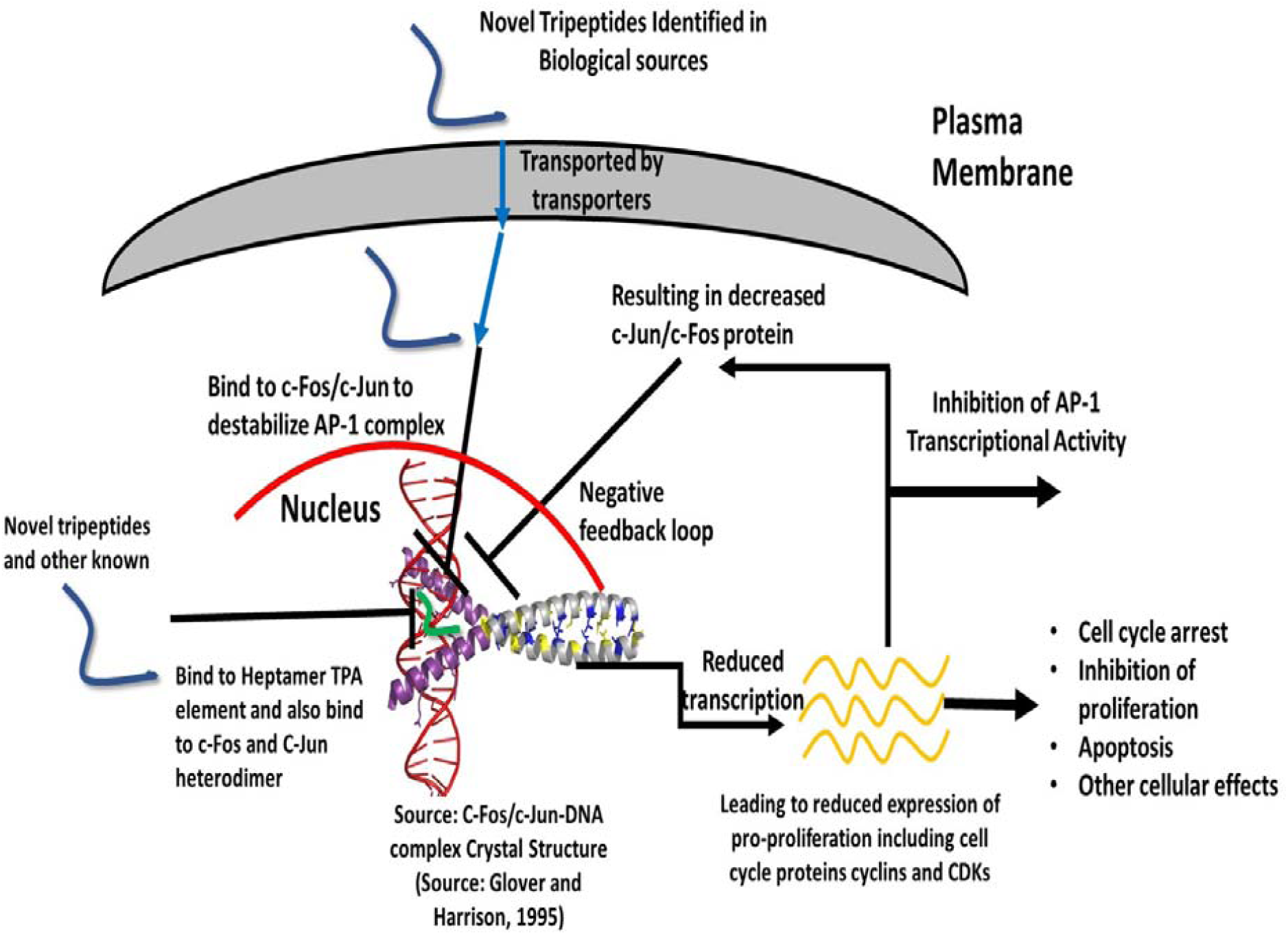
This flow diagram proposes avenues for chemical inhibition of AP-1 transcriptional complex and use of GUDF derived peptides as potential anticancer agents. The strong binding affinity and specific docking within the AP-1 transcriptional complex are displayed by these tripeptides. Due to these properties, selected tripeptides are suggested to show the inhibition of proliferation of cancer cells, since disruption of AP-1 transcriptional complex is linked with the cell cycle arrest and blocked for proliferation.

## FUTURE DIRECTIONS AND CONCLUSION

In conclusion, findings suggest the potential inhibitory role of novel tripeptides such as Glu-Glu-Arg, Gly-Arg-Pro and Gln-Lys-Arg upon the AP-1 transcriptional complex based on the in silico studies. Interestingly, these tripeptides including Glu-Glu-Arg, Gly-Arg-Pro and Gln-Lys-Arg are present in the intracellular compartment of HCT-116 cells treated by GUDF enriched with tripeptides. Further, these data suggested that GUDF enriched with tripeptides treatment to HCT-116 cells resulted in a clear proliferation arrest. Based on the existing views, inhibition of AP-1 complex leads to similar proliferation arrest in cancer cells. Therefore, authors project these tripeptides including Glu-Glu-Arg, Gly-Arg-Pro and Gln-Lys-Arg as potential candidates that may block AP-1 mediated oncogene expressions and in turn will retard the growth and proliferation of cancer cells. The authors see some limitation of present findings on the lack of in vitro experiments on actual binding of these tripeptides with the heptamer consensus sequence of AP-1 complex. These encouraging findings warrants future investigations at in vitro, preclinical and clinical levels to evaluate their role as antiproliferative agents. In future, a possibility of turning these tripeptides into self-assembled nano-particles after conjugation with metal ions is proposed for its better delivery into the targeted cancer cells.

## ACKNOWLEDGEMENTS

The authors acknowledge financial support from DST-SERB, Government of India, New Delhi, India (SERB/LS-1028/2013) and Dr. D.Y. Patil Vidyapeeth, Pune, India (DPU/05/01/2016).

## DETAILS OF SUPPLEMENTARY FIGURES

**Figure S1.**
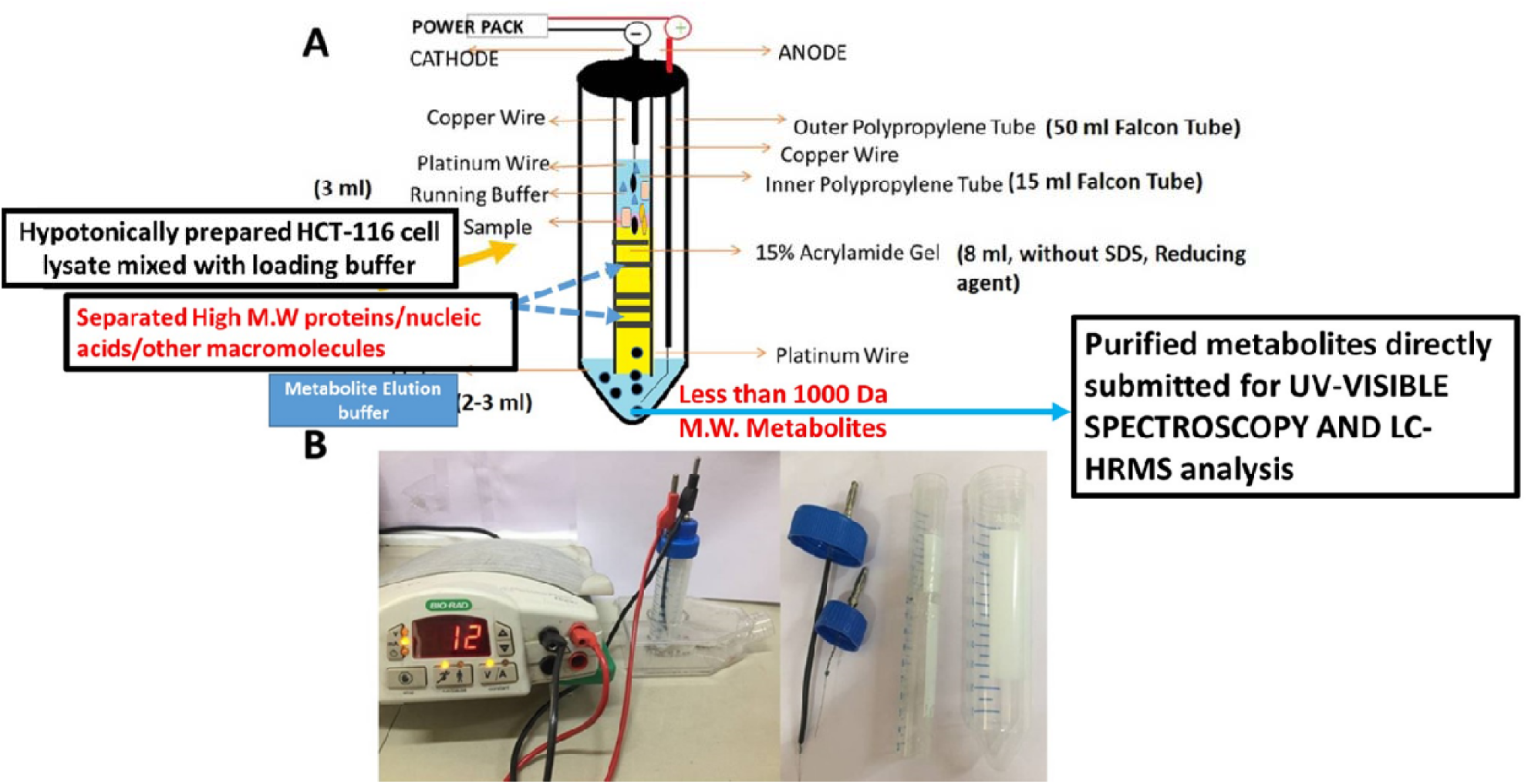
A design and working model of a novel VTGE metabolite purification system from biological materials such as cell lysates, tissue lysates, biological fluids and nails lysate. **(A)** An assembly and design of VTGE system is illustrated. **(B)** A working model is depicted for VTGE system.

**Figure S2:**
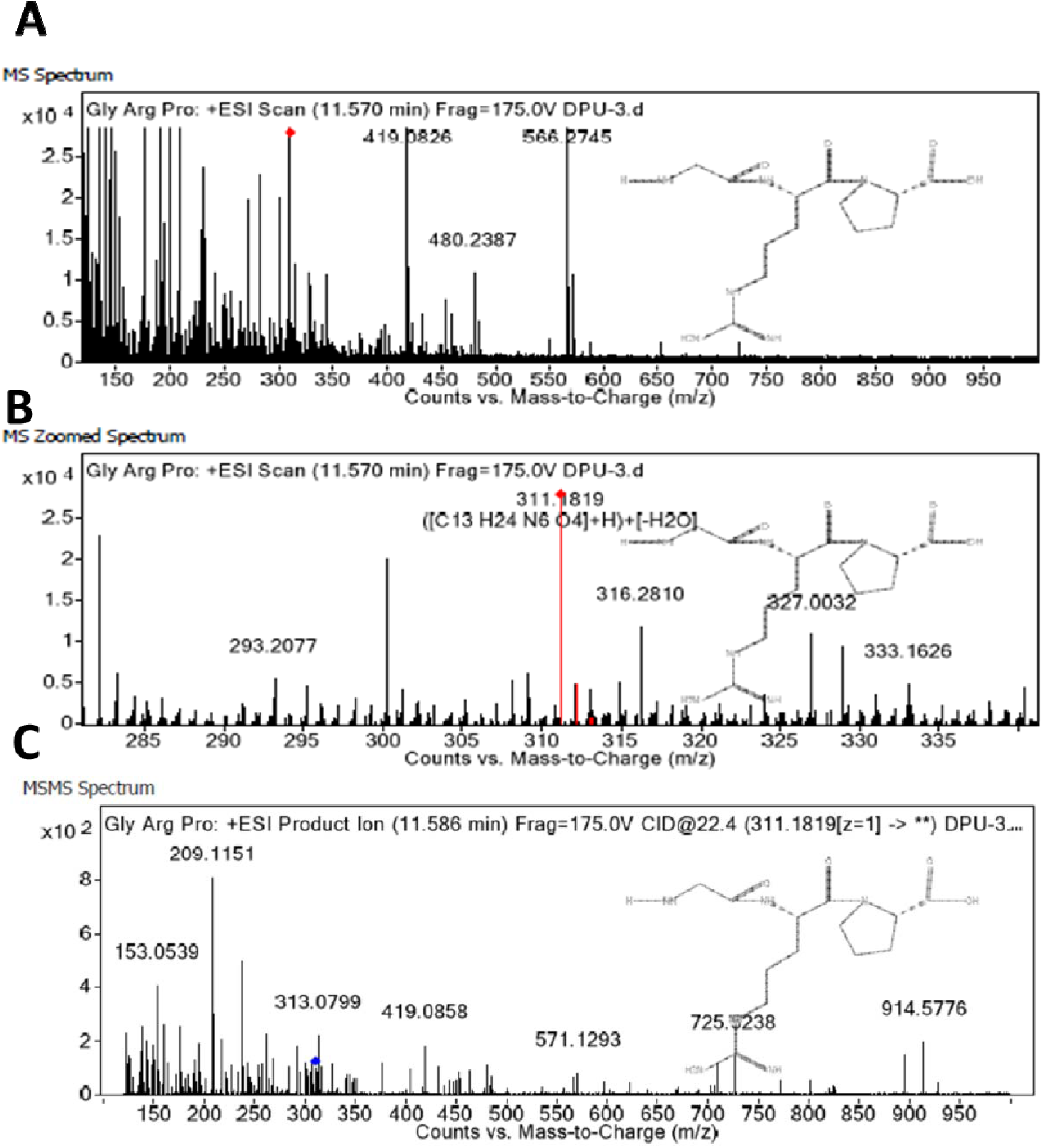
A LC-HRMS mass ion spectra of tripeptide Gly-Arg-Pro identified as intracellular tripeptides in GUDF treated HCT-116 cells assisted by VTGE metabolite purification system. Hypotonically prepared cell lysate of HCT-116 cells treated with GUDF was purified by employing a novel methodology VTGE and analyzed by LC-HRMS technique in positive ESI-MS mode. (A) LC-HRMS positive ESI-MS spectra of Gly-Arg-Pro. (B) A zoomed MS ion spectra of tripeptide Gly-Arg-Pro. (C) A MS/MS positive ESI spectra with product mass ion 153.0539, 313.0539, 419.0858, 571.1291, 725.5238, 914.5776 of Gly-Arg-Pro.

**Figure S3:**
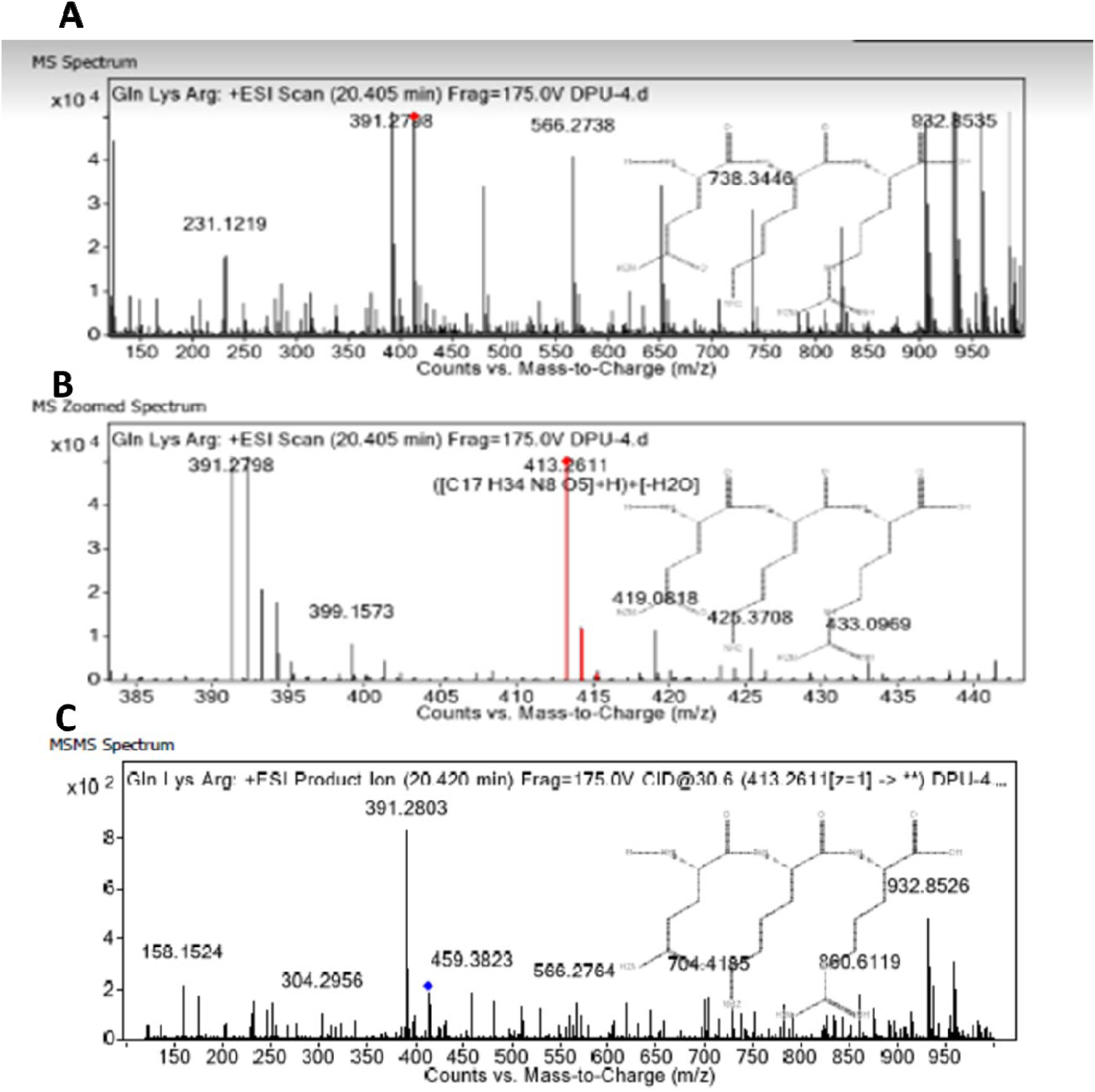
A LC-HRMS mass ion spectra of tripeptide Gln-Lys-Arg identified as intracellular tripeptides in GUDF treated HCT-116 cells assisted by VTGE metabolite purification system. Hypotonically prepared cell lysate of HCT-116 cells treated with GUDF was purified by employing a novel methodology VTGE and analyzed by LC-HRMS technique in positive ESI-MS mode. (A) LC-HRMS positive ESI-MS spectra of Gln-Lys-Arg. (B) A zoomed MS ion spectra of tripeptide Gln-Lys-Arg. (C) A MS/MS positive ESI spectra with product mass ion (158.1524, 304.2956, 391.2803, 459.3823, 566.2764, 704.4185, 860.8119, 932.8526) of Gln-Lys-Arg.

**Figure S4:**
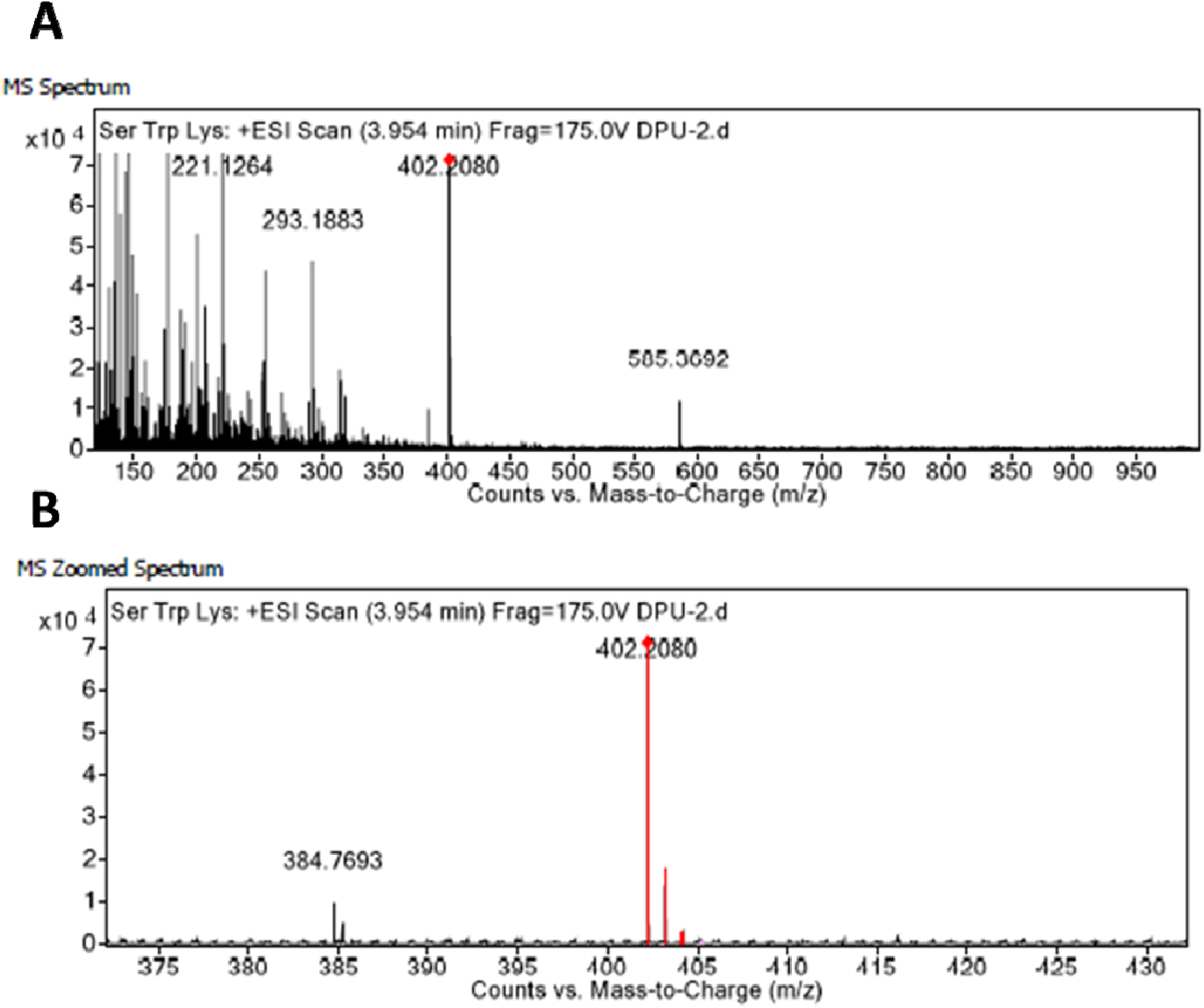
A LC-HRMS mass ion spectra of tripeptide Ser-Trp-Lys identified as intracellular tripeptides in GUDF treated HCT-116 cells assisted by VTGE metabolite purification system. Hypotonically prepared cell lysate of HCT-116 cells treated with DMSO was purified by employing a novel methodology VTGE and analyzed by LC-HRMS technique in positive ESI-MS mode. (A) LC-HRMS positive ESI-MS spectra of Ser-Trp-Lys. (B) A zoomed MS ion spectra of tripeptide Ser-Trp-Lys.

**Figure S5:**
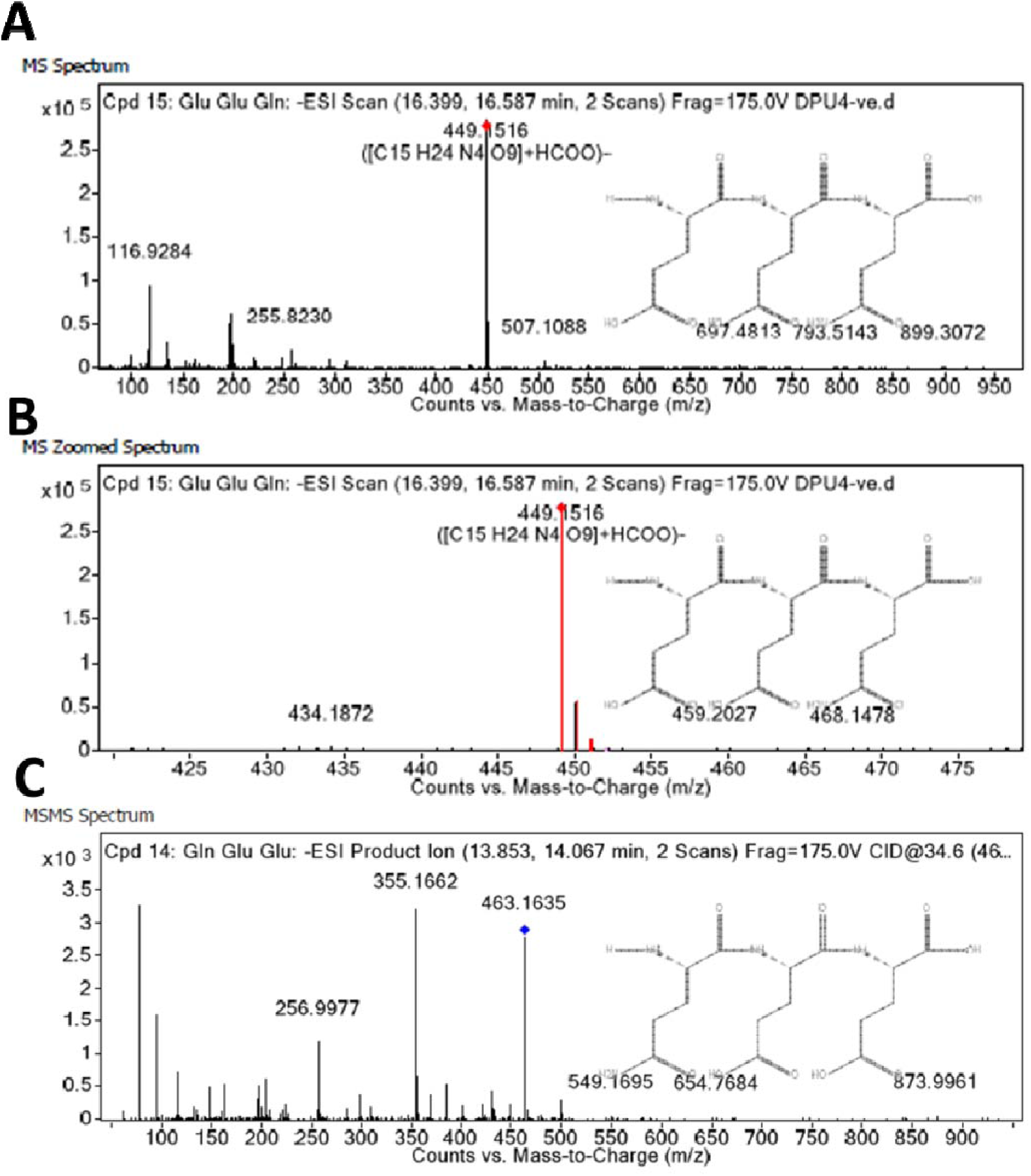
A LC-HRMS mass ion spectra of tripeptide Glu-Glu-Gln identified as intracellular tripeptides in DMSO treated HCT-116 cells assisted by VTGE metabolite purification system. Hypotonically prepared cell lysate of HCT-116 cells treated with GUDF was purified by employing a novel methodology VTGE and analyzed by LC-HRMS technique in positive ESI-MS mode. (A) LC-HRMS positive ESI-MS spectra of Glu-Glu-Gln. (B) A zoomed MS ion spectra of tripeptide Glu-Glu-Gln. (C) A MS/MS positive ESI spectra with product mass ion (256.9977, 355.1662, 463.1635, 549.1695, 654.7684, 873.9961) of Glu-Glu-Gln.

**Figure S6:**
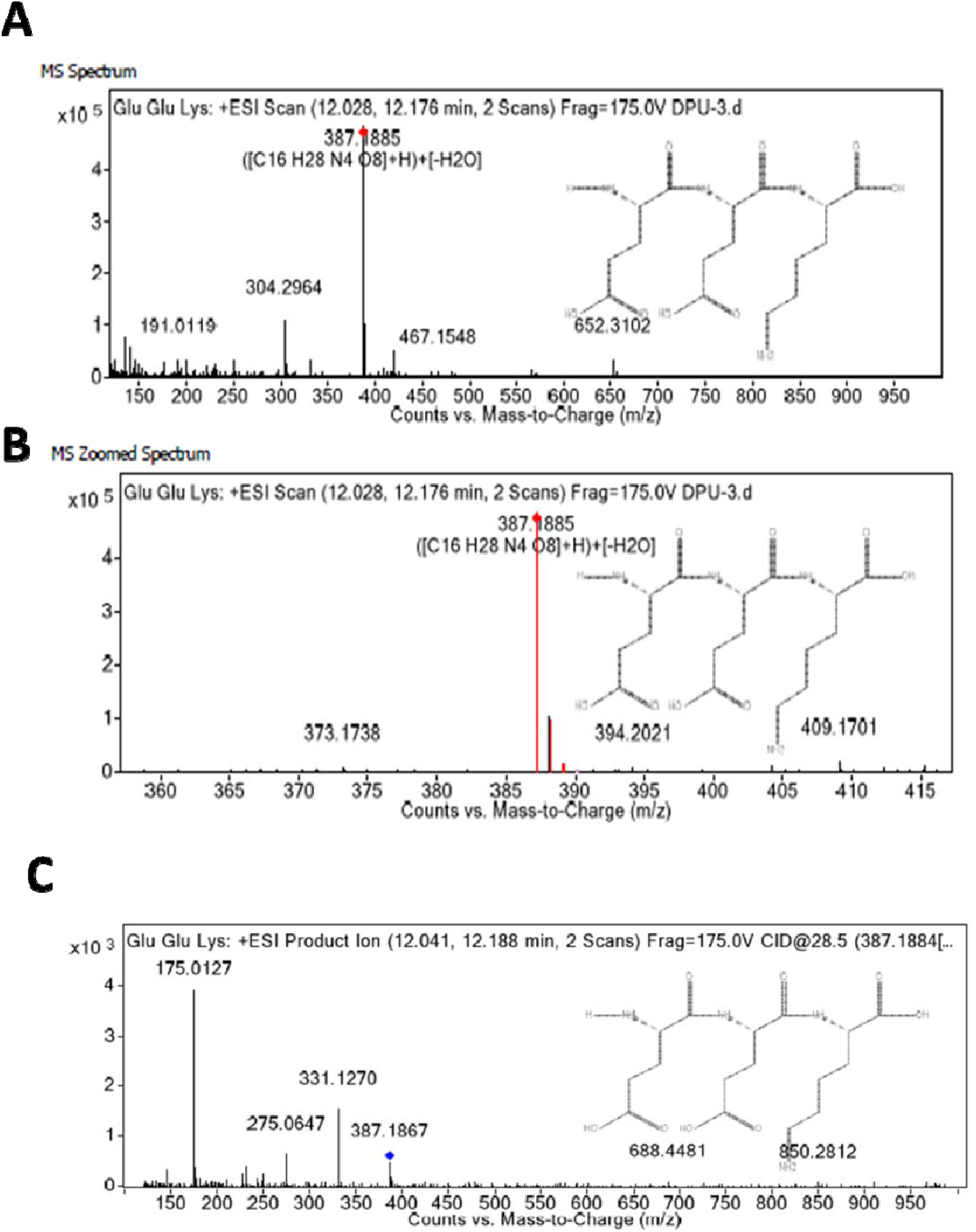
A LC-HRMS mass ion spectra of tripeptide Glu-Glu-Lys identified as intracellular tripeptides in both DMSO and GUDF treated HCT-116 cells assisted by VTGE metabolite purification system. Hypotonically prepared cell lysate of HCT-116 cells treated with GUDF was purified by employing a novel methodology VTGE and analyzed by LC-HRMS technique in positive ESI-MS mode. (A) LC-HRMS positive ESI-MS spectra of Glu-Glu-Lys. (B) A zoomed MS ion spectra of tripeptide Glu-Glu-Lys. (C) A MS/MS positive ESI spectra with product mass ion (275.0647, 331.1270, 387.1867, 688.4481, 850.2812) of Glu-Glu-Lys.

**Figure S7:**
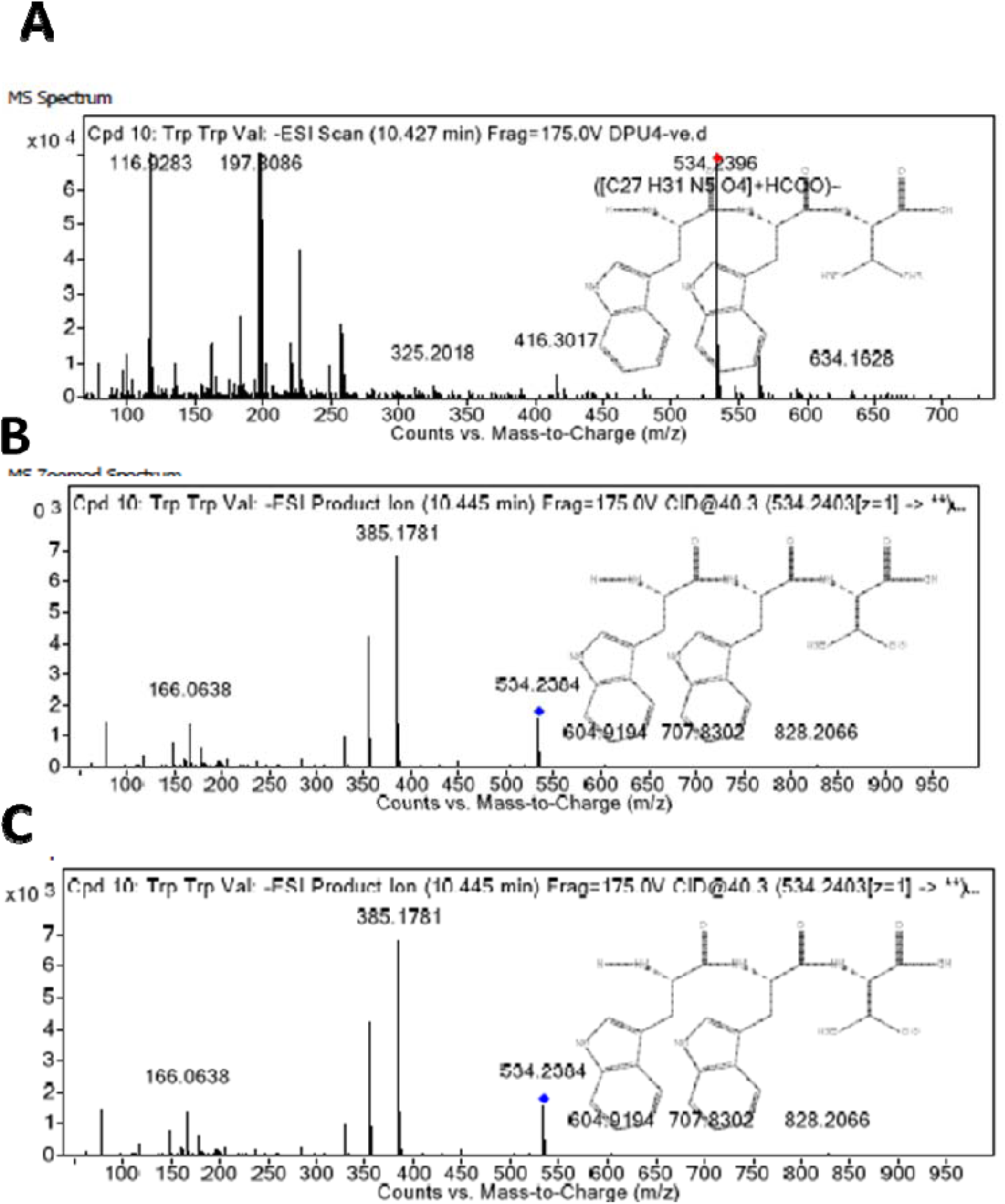
A LC-HRMS mass ion spectra of tripeptide Trp-Trp-Val identified as intracellular tripeptides in GUDF treated HCT-116 cells assisted by VTGE metabolite purification system. (166.0638, 385.1781, 534.2064, 604.9194, 707.8302, 828.2066) Hypotonically prepared cell lysate of HCT-116 cells treated with GUDF was purified by employing a novel methodology VTGE and analyzed by LC-HRMS technique in negative ESI-MS mode. (A) LC-HRMS negative ESI-MS spectra of Trp-Trp-Val. (B) A zoomed MS ion spectra of tripeptide Trp-Trp-Val. (C) A MS/MS negative ESI spectra with product mass ion (166.0638, 385.1781, 534.2064, 604.9194, 707.8302, 828.2066) of Trp-Trp-Val.

**Figure S8:**
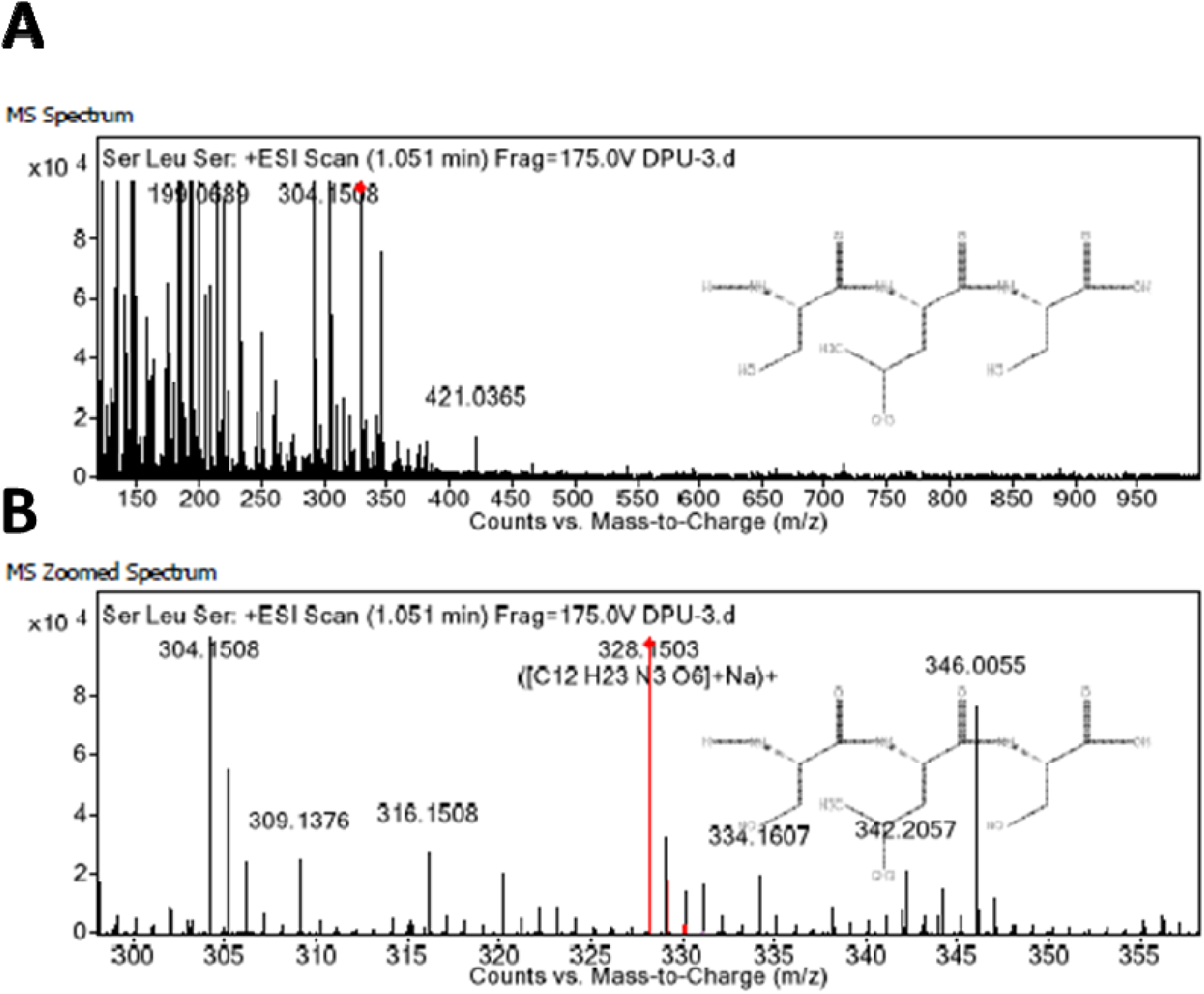
A LC-HRMS mass ion spectra of tripeptide Ser-Leu-Ser identified as intracellular tripeptides in DMSO treated HCT-116 cells assisted by VTGE metabolite purification system. Hypotonically prepared cell lysate of HCT-116 cells treated with GUDF was purified by employing a novel methodology VTGE and analyzed by LC-HRMS technique in positive ESI-MS mode. (A) LC-HRMS positive ESI-MS spectra of Ser-Leu-Ser. (B) A zoomed MS ion spectra of tripeptide Ser-Leu-Ser.

**Figure S9.**
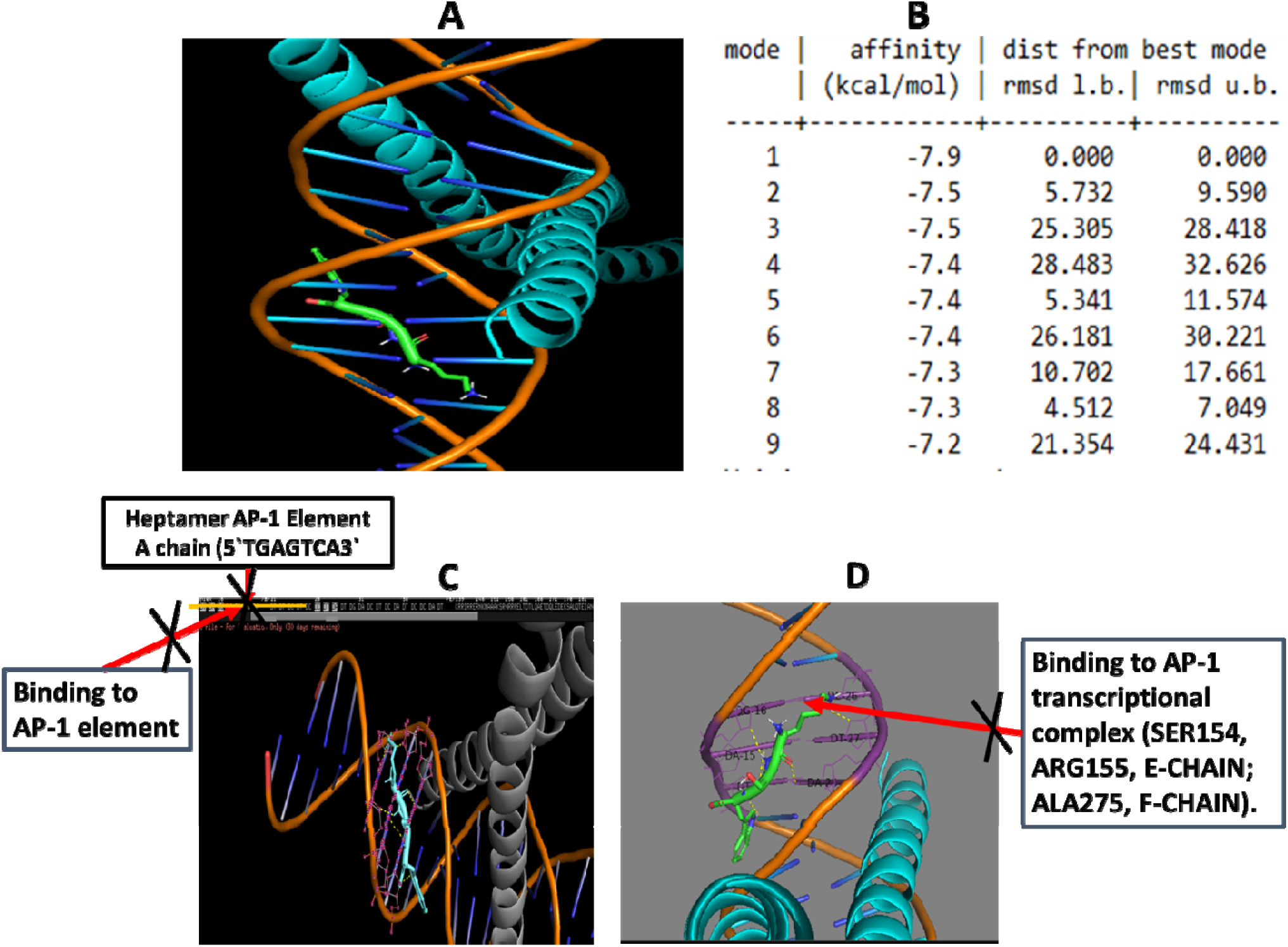
An intracellular tripeptide Lys-Ser-Trp from DMSO treated HCT-116 cells does not show a specific binding to AP-1 transcriptional complex and low docking energy value is at −6.0 Kcal/Mol. (A). A PyMol view of docking between tripeptide Lys-Ser-Trp and c-Jun:c-Fos:DNA ternary complex. (B) Value on docking affinity obtained during AutoDock Vina based molecular interaction between tripeptide Ser-Leu-Ser and c-Jun:c-Fos:DNA ternary complex. (C). In depth image of PyMol generated model shows that tripeptide Lys-Ser-Trp is not able to bind within the heptamer AP-1 consensus site (5′-TGAGTCA-3′) of AP-1 transcriptional complex. (D) Also, tripeptide Lys-Ser-Trp does not bind to E-chain (c-Fos) and F-chain (c-Jun) heterodimer complex within the AP-1 transcriptional complex.

**Figure S10.**
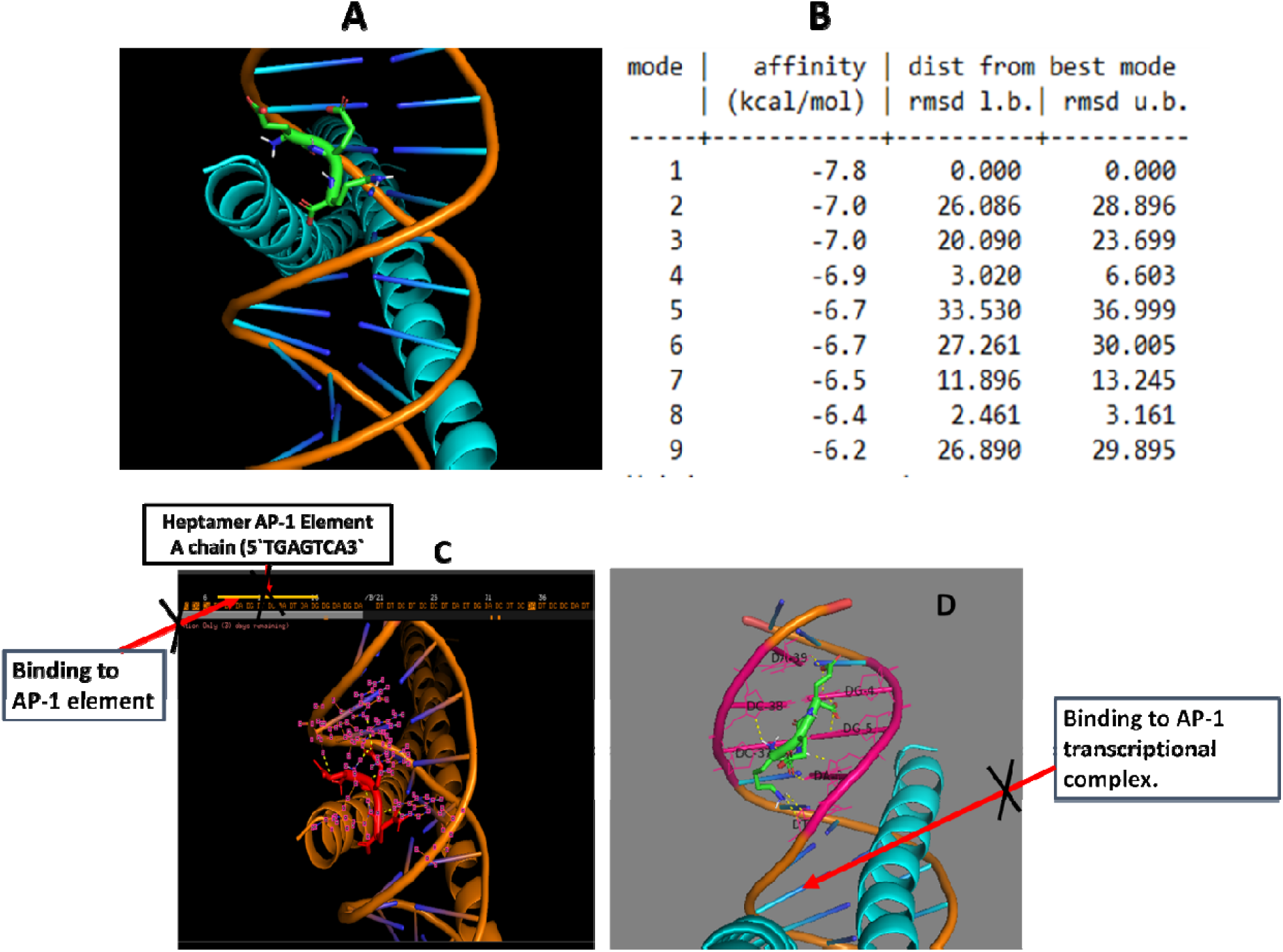
An intracellular tripeptide Glu-Glu-Gln from DMSO treated HCT-116 cells does not show a specific binding to AP-1 transcriptional complex and low docking energy value is at −7.8 Kcal/Mol. (A). A PyMol view of docking between tripeptide Glu-Glu-Gln and c-Jun:c-Fos:DNA ternary complex. (B) Value on docking affinity obtained during AutoDock Vina based molecular interaction between tripeptide Ser-Leu-Ser and c-Jun:c-Fos:DNA ternary complex. (C). In depth image of PyMol generated model shows that tripeptide Glu-Glu-Gln is not able to bind within the heptamer AP-1 consensus site (5′-TGAGTCA-3′) of AP-1 transcriptional complex. (D) Also, tripeptide Glu-Glu-Gln does not bind to E-chain (c-Fos) and F-chain (c-Jun) heterodimer complex within the AP-1 transcriptional complex.

**Figure S11.**
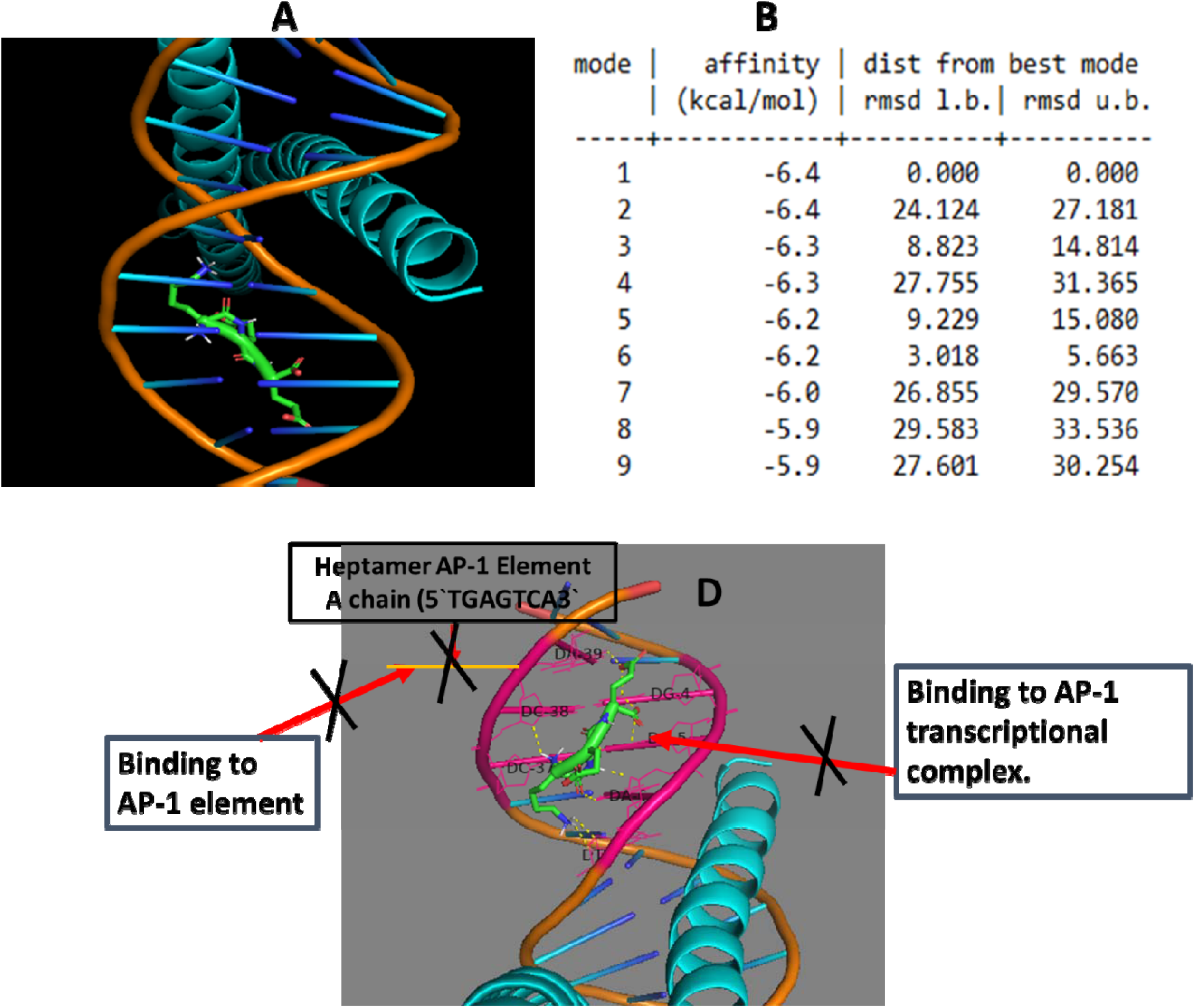
An intracellular tripeptide Glu-Glu-Lys from DMSO and GUDF treated HCT-116 cells does not show a specific binding to AP-1 transcriptional complex and low docking energy value is at −6.4 Kcal/Mol. (A). A PyMol view of docking between tripeptide Glu-Glu-Lys and c-Jun:c-Fos:DNA ternary complex. (B) Value on docking affinity obtained during AutoDock Vina based molecular interaction between tripeptide Glu-Glu-Lys and c-Jun:c-Fos:DNA ternary complex. (C). In depth image of PyMol generated model shows that tripeptide Glu-Glu-Lys is not able to bind within the heptamer AP-1 consensus site (5′-TGAGTCA-3′) of AP-1 transcriptional complex. Also, tripeptide Glu-Glu-Lys does not bind to E-chain (c-Fos) and F-chain (c-Jun) heterodimer complex within the AP-1 transcriptional complex.

**Figure S12.**
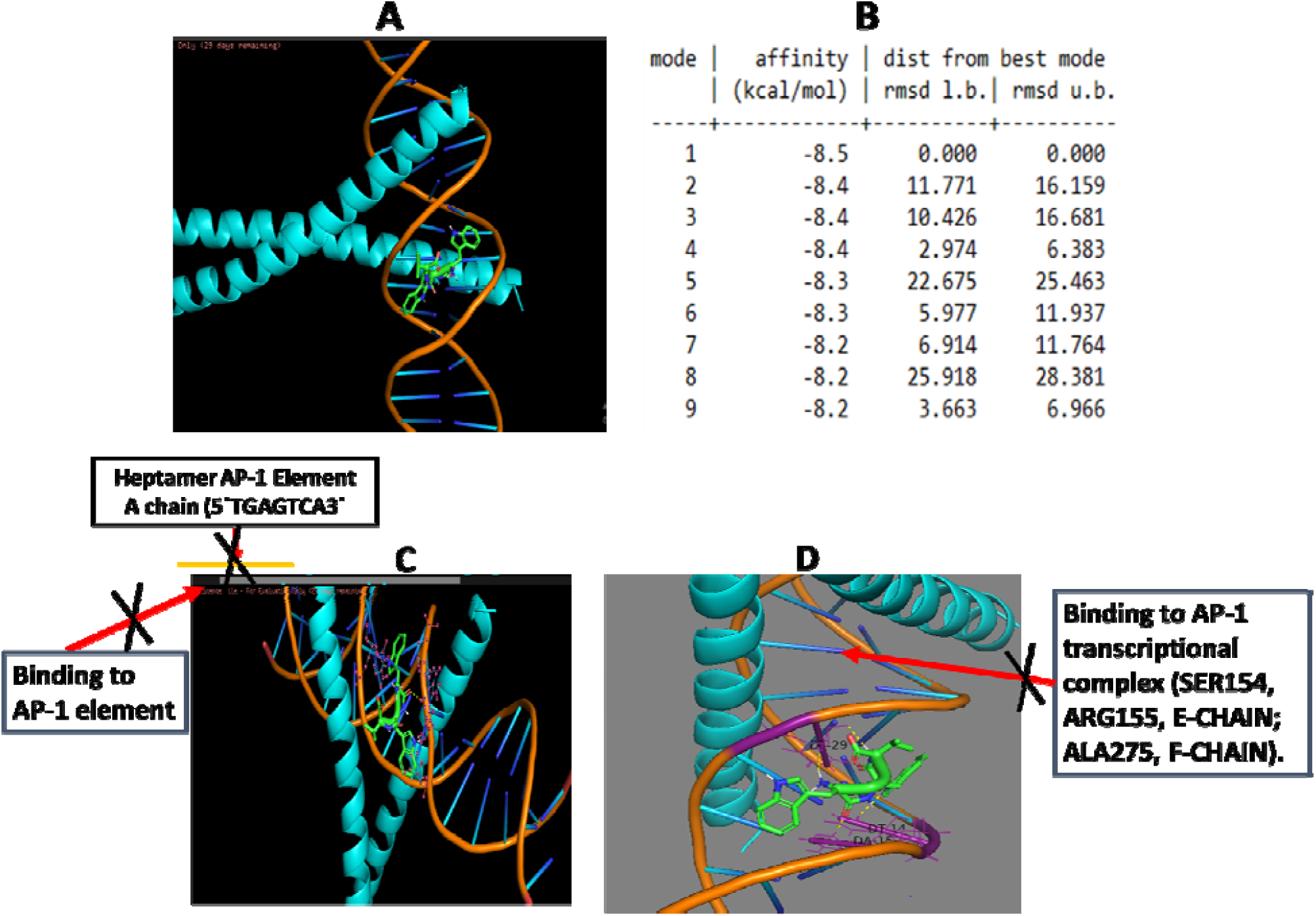
An intracellular tripeptide Trp-Trp-Val from GUDF treated HCT-116 cells does not show a specific binding to AP-1 transcriptional complex and low docking energy value is at −8.5 Kcal/Mol. (A). A PyMol view of docking between tripeptide Trp-Trp-Val and c-Jun:c-Fos:DNA ternary complex. (B) Value on docking affinity obtained during AutoDock Vina based molecular interaction between tripeptide Trp-Trp-Val and c-Jun:c-Fos:DNA ternary complex. (C). In depth image of PyMol generated model shows that tripeptide Trp-Trp-Val is not able to bind within the heptamer AP-1 consensus site (5′-TGAGTCA-3′) of AP-1 transcriptional complex. (D) Also, emphasized image indicates that tripeptide Trp-Trp-Val is not bound to E-chain (c-Fos) and F-chain (c-Jun) heterodimer complex within the AP-1 transcriptional complex.

**Figure S13.**
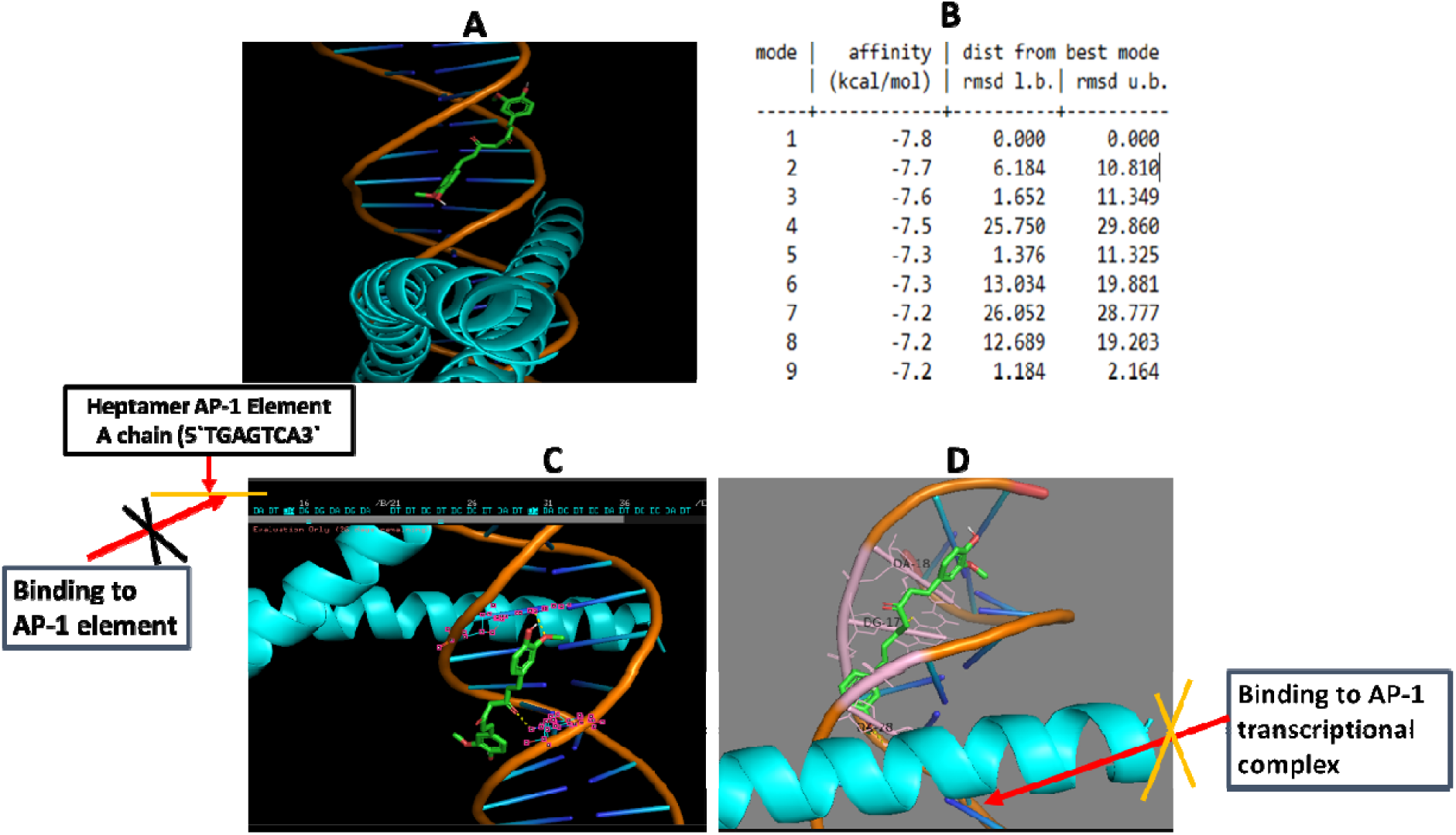
As a reference well-known DNA binding and anticancer agent curcumin does not show a specific binding to AP-1 transcriptional complex and with a docking energy value is at - 7.8 Kcal/Mol. (A). A PyMol view of docking between curcumin and c-Jun:c-Fos:DNA ternary complex. (B) Value on docking affinity obtained during AutoDock Vina based molecular interaction between curcumin and c-Jun:c-Fos:DNA ternary complex. (C). In depth image of PyMol generated model shows that curcumin is not able to bind within the heptamer AP-1 consensus site (5′-TGAGTCA-3′) of AP-1 transcriptional complex. (D) Also, emphasized image indicates that curcumin is not bound to E-chain (c-Fos) and F-chain (c-Jun) heterodimer complex within the AP-1 transcriptional complex.

**Figure S14.**
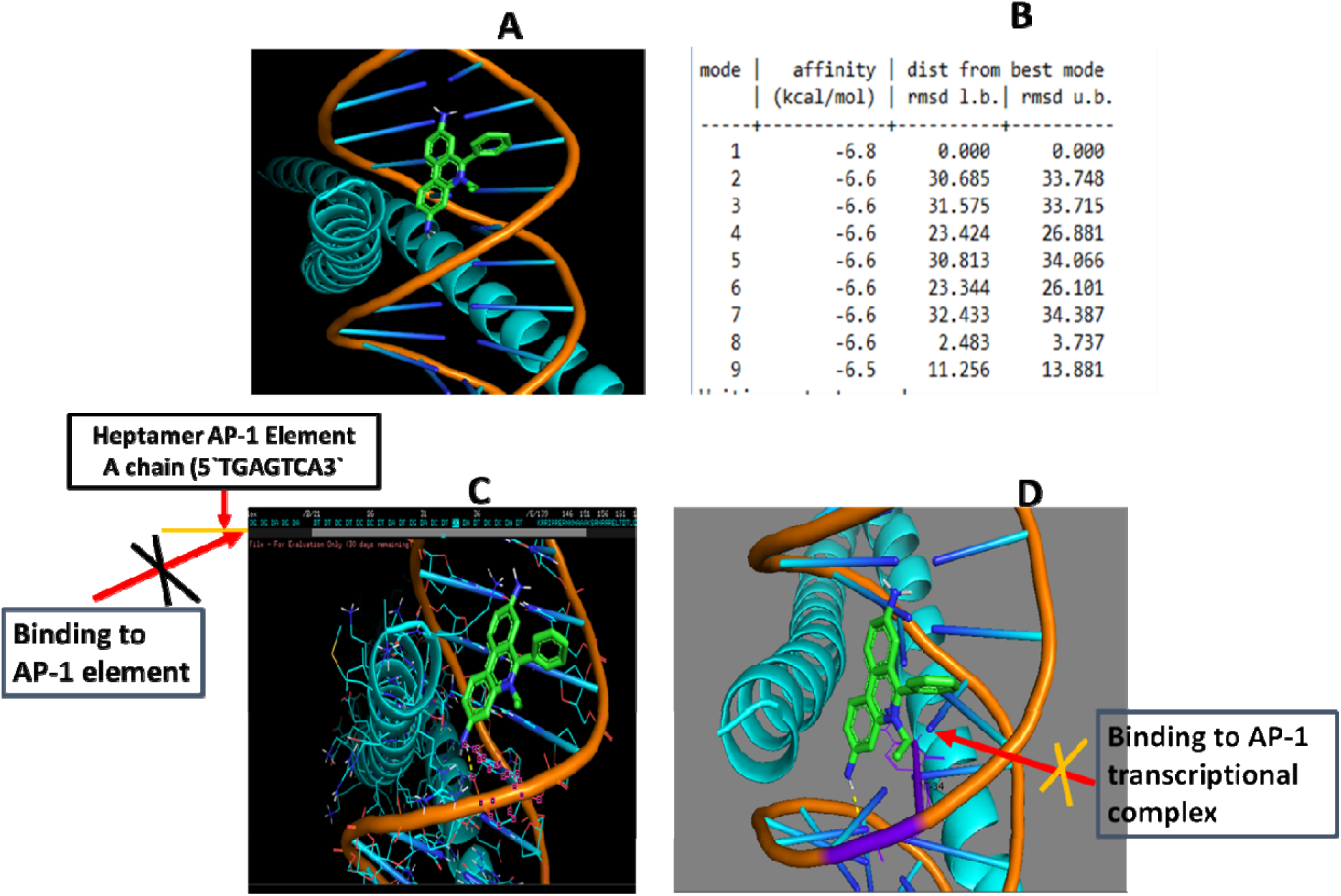
As a reference, well-known DNA intercalating agent ethidium bromide does not show a specific binding to AP-1 transcriptional complex and non-specific docking energy value is at −6.8 Kcal/Mol. (A). A PyMol view of docking between ethidium bromide and c-Jun:c-Fos:DNA ternary complex. (B) Value on docking affinity obtained during AutoDock Vina based molecular interaction between ethidium bromide and c-Jun:c-Fos:DNA ternary complex. (C). In depth image of PyMol generated model shows that ethidium bromide is not able to bind within the heptamer AP-1 consensus site (5′-TGAGTCA-3′) of AP-1 transcriptional complex. (D) Also, emphasized image indicates that ethidium bromide is not bound to E-chain (c-Fos) and F-chain (c-Jun) heterodimer complex within the AP-1 transcriptional complex.

**Figure S15.**
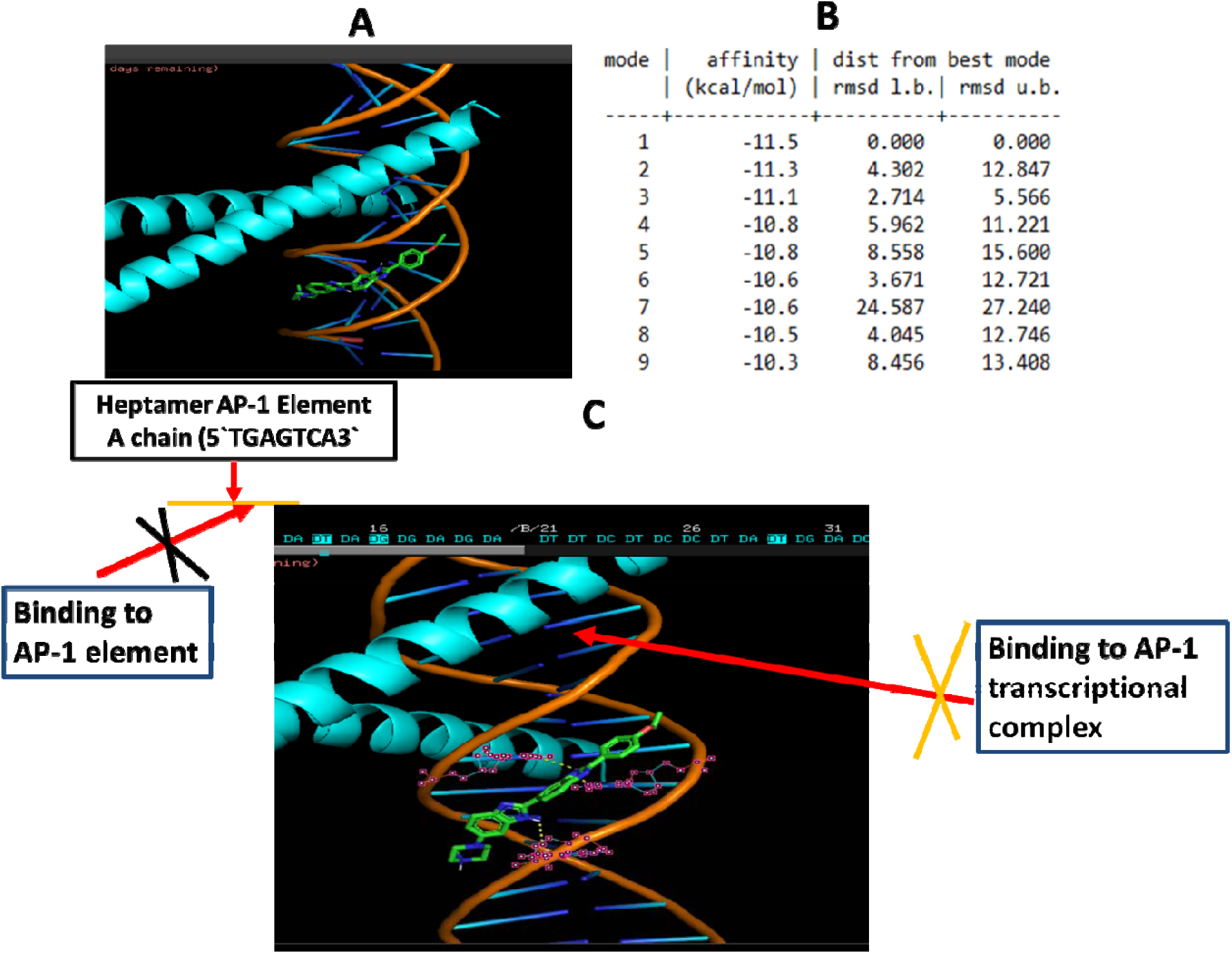
As a reference, well-known DNA binding dye Hoechst 33342 does not show a specific binding to AP-1 transcriptional complex and however, non-specific DNA binding is high with a docking energy value is at −11.5 Kcal/Mol. (A). A PyMol view of docking between Hoechst 33342 and c-Jun:c-Fos:DNA ternary complex. Value on docking affinity obtained during AutoDock Vina based molecular interaction between Hoechst 33342 and c-Jun:c-Fos:DNA ternary complex. (C). In depth image of PyMol generated model shows that Hoechst 33342 is not able to bind within the heptamer AP-1 consensus site (5′-TGAGTCA-3′) of AP-1 transcriptional complex. (D) Also, emphasized image indicates that Hoechst 33342 is not bound to E-chain (c-Fos) and F-chain (c-Jun) heterodimer complex within the AP-1 transcriptional complex.

